# Mapping the THESEUS1 interactome links cell wall integrity and stress signalling in Arabidopsis

**DOI:** 10.64898/2026.10.22.760881

**Authors:** Steven T. W. Zwartkruis, Wiebke Haeger, Tereza Tichá, Nelly Braun, Zdenka Bartosova, Animesh Sharma, Adam Zeiner, Gregor Madej, Vivien J. Klein, Dhika Amanda, Luis Alonso Baez, Elke Barbez, Michael Wrzaczek, Lars Hagen, Thorsten Hamann

**Author notes:** Corresponding authors: Thorsten Hamann, Department of Biology, Faculty of Natural Sciences, Norwegian University of Science and Technology, Trondheim, Norway,. Steven T. W. Zwartkruis, Department of Biology, Faculty of Natural Sciences, Norwegian University of Science and Technology, Trondheim, Norway. The author(s) responsible for distribution of materials integral to the findings presented in this article in accordance with the policy described in the Author guidelines: Thorsten Hamann.

## Abstract

The receptor kinase THESEUS1 (THE1) activates responses to impairment of cell wall integrity (CWI). However, the interaction partners enabling THE1 to mediate the responses remain underexplored, impeding a mechanistic understanding of THE1. We produced a set of THE1-GFP-fusion constructs in which THE1 is either wild type, mutated in its kinase domain, has a loss-of-function mutation in its ectodomain, or lacks the kinase or extracellular domain. We used these constructs to perform localization studies and immunoprecipitation-liquid chromatography-tandem mass spectrometry under control and stress conditions (isoxaben-induced cell wall damage and sorbitol-induced hyperosmotic stress) to identify the THE1 interactome. The interactome of THE1 changed in a context-dependent manner, linking THE1 to known signaling components in CWI maintenance and phytohormone response (such as proton pumps, calcium pumps, other receptor kinases or cellulose synthesis regulators). Moreover, we identified novel interactors, associating THE1 with hitherto unexplored signaling pathways in response to CWI impairment. Our results also suggest that the THE1_G37D_ protein (corresponding to the *the1-1* allele) is retained in the endoplasmic reticulum, explaining its loss-of-function phenotype. The THE1 interactome and localization data provide the foundation to build a mechanistic understanding of THE1-mediated CWI maintenance.

## Introduction

The plant cell wall is a multifunctional structure that regulates cell shape, influences plant development, and forms the primary barrier protecting plant cells against environmental stresses (Novaković et al. 2018; Cosgrove 2024b; Munzert and Engelsdorf 2025; Perez-Alonso et al. 2025). Composition and structural coherence determine the integrity of the plant cell wall and its mechanical properties (Cosgrove 2024b). Maintaining cell wall integrity (CWI) is crucial to plants as plant cells represent pressurized structures which swell or burst if not resisted by a functional cell wall (Wolf 2022). While coordinated loosening of cell walls is required for turgor-driven expansion during growth, CWI impairment can be a sign of biotic or abiotic stress such as pathogen attack or drought (Munzert and Engelsdorf 2025; Debnath et al. 2026). To maintain the cell wall barrier during plant development and in response to environmental stress, plants evolved a CWI maintenance mechanism (Rui and Dinneny 2020; Alonso Baez et al. 2022; Wolf 2022). Plasma membrane-localized mechanosensitive ion channels and receptor kinases (RKs) are molecular components of the CWI maintenance signaling pathways (Alonso Baez et al. 2022; Wolf 2022). Mechanosensitive ion channels allow ion flux upon changes in plasma membrane tension following cell wall weakening or changes in turgor pressure (Bacete and Hamann 2020; Guichard et al. 2022). RKs monitor cell wall state by sensing extracellularly localized ligands such as cell wall-derived carbohydrates or small peptides (Tseng et al. 2022; Lu et al. 2025; Debnath et al. 2026). The *Catharanthus roseus* RECEPTOR-LIKE KINASE 1-LIKE (*Cr*RLK1L) RK family plays a central role in CWI maintenance (Lindner et al. 2012; Alonso Baez et al. 2022; Manchanda and Geitmann 2025; Schoenaers and Vissenberg 2025). The ectodomain of these receptors has been proposed to sense CWI through interactions with RAPID ALKALINIZATION FACTOR (RALF) peptides and the cell wall polysaccharide pectin (Manchanda and Geitmann 2025; Schoenaers and Vissenberg 2025). *Arabidopsis thaliana* plants with mutations in *CrRLK1L*s display altered development and plant stress responses (Duan et al. 2010; Engelsdorf et al. 2018; Schoenaers et al. 2018; Zhao et al. 2021; Gigli-Bisceglia et al. 2022; Leicher et al. 2026). For example, root hairs in loss-of-function mutants of the *Cr*RLK1L genes *FERONIA* (*FER*) and *ERULUS* burst or bulge (Duan et al. 2010; Bai et al. 2014; Schoenaers et al. 2018), indicating impaired coordination between turgor pressure and CWI. Moreover, *fer-4* loss-of-function seedlings display constitutive activation of the jasmonic acid (JA) and abscisic acid (ABA) phytohormone pathways (Guo et al. 2018; Wang et al. 2022). Thus, studying these *Cr*RLK1Ls is important to understand how CWI is monitored and linked to stress signaling pathways.

The *Cr*RLK1L family member THESEUS1 (THE1) was the first RK to be associated with CWI maintenance in *A. thaliana* (Hématy et al. 2007; Zwartkruis et al. 2026). THE1 consists of a signal peptide, two extracellular malectin-like domains, a single transmembrane helix and a cytosolic kinase domain (Hématy et al. 2007; Zwartkruis et al. 2026). THE1 was initially discovered in a mutant-suppressor screen of the *CELLULOSE SYNTHASE 6* mutant *procuste1-1* (*prc1-1)* in *A. thaliana* (Hématy et al. 2007). Plants homozygous for the *the1-1* loss-of-function allele (containing a G37D mutation in the first malectin-like domain) suppress a hypocotyl elongation defect in etiolated *prc1-1* seedlings (Hématy et al. 2007). THE1 is also responsible for ectopic lignification and immune system activation induced by isoxaben (ISX), a cellulose biosynthesis inhibitor impairing CWI (Scheible et al. 2001; Engelsdorf et al. 2018; Gonneau et al. 2018; Larson and McFarlane 2021; Zhai et al. 2024). The requirement of THE1 in activating responses to cellulose biosynthesis inhibition led to its proposed role in CWI maintenance (Hématy et al. 2007). Moreover, THE1 influences the production of the plant stress hormones JA and ABA. When CWI is challenged by ISX treatment, *the1-1* mutant seedlings produce less JA than wild-type *A. thaliana* seedlings (Engelsdorf et al. 2018; Bacete et al. 2022). More recently, *the1-1* mutants were found to produce more ABA after treatment with the osmoticum sorbitol (Bacete et al. 2022). Therefore, THE1 is considered a positive regulator of JA and a negative regulator of ABA production. The T-DNA insertion in the hypermorphic *the1-4* mutant leads to a THE1 version truncated 32 amino acid residues after the transmembrane helix, hence lacking the entire kinase domain (Guo et al. 2009; Merz et al. 2017). This mutation results in stronger ISX-induced responses than observed in wild-type plants (Merz et al. 2017; Engelsdorf et al. 2018; Bacete et al. 2022). It is not fully understood how *the1-4* lacking a functional THE1 kinase domain activates enhanced cellulose biosynthesis inhibition responses. It has been suggested that the THE1 ectodomain or transmembrane domain could act through heteromeric complexes with other RKs to elicit such responses, and that the THE1 kinase could be part of a negative feedback loop (Merz et al. 2017). However, the components of these complexes as well as downstream signaling components remain to be determined.

To date, only few interaction partners of THE1 have been described. As a THE1 ligand, the cysteine-rich peptide RALF34 induces THE1-dependent apoplast alkalinization and calcium influx (Gonneau et al. 2018). GUANINE EXCHANGE FACTOR 4 (GEF4) was described to interact with THE1 and positively regulate defense responses against the fungal pathogen *Botrytis cinerea* (Qu et al. 2017). Recently, the ectodomain of THE1 was described to be part of the receptor complex that establishes polytubey block, based on co-immunoprecipitation (co-IP) of the THE1 ectodomain with the ectodomains of the *Cr*RLK1Ls FER, ANJEA (ANJ), and HERCULES RECEPTOR KINASE 1 (HERK1), and RALF6, -7, -16, -36, and -37 (Li et al. 2026a). *Cr*RLK1L heteromerization involving multiple *Cr*RLK1Ls and multiple RALFs are hypothesized to determine specific signaling outputs (Schoenaers and Vissenberg 2025). In comparison to other RKs (Cheung 2024; Delesalle et al. 2024), the number of interactors described for THE1 is low (Zwartkruis et al. 2026). More knowledge about the interactors of THE1 is needed to understand its mode of action and the signaling pathways it is involved in during CWI maintenance.

Here, we studied the interactome of THE1 by immunoprecipitation-liquid chromatography-tandem mass spectrometry (IP-LC-MS/MS). Previously, this method has been used to identify interactors of other RKs such as BRASSINOSTEROID INSENSITIVE 1 (BRI1) (Bai et al. 2026) and FER (Leicher et al. 2026). We identified novel THE1 interactors including other RKs, ubiquitin ligases, proton pumps, calcium pumps, ion channels, nuclear transport proteins, and proteins associated with the regulation of cellulose biosynthesis. We functionally analyzed THE1-GFP transgenic plants in which domains of THE1 are mutated or deleted by measuring JA levels, visualizing subcellular localization, and identifying their interactome. Our results showed that the THE1_G37D_ protein, encoded in *the1-1*, apparently mislocalizes to the endoplasmic reticulum (ER) and interacts with ER-localized molecular chaperones. Overall, our results provide new insights into the molecular pathways by which THE1 links CWI to plant stress responses.

## Materials and methods

### Plant material and growth conditions

The *A. thaliana* lines *the1-1* (Hématy et al. 2007), *the1-4* (Guo et al. 2009), *the1-6* (Merz et al. 2017), and the line expressing *35S_pro_:LTI6B-GFP* (Cutler et al. 2000) in Col-0 background were obtained from the laboratories previously publishing them. Seeds were surface sterilized (5 min 70% (v/v) ethanol, 5 min 50% (v/v) bleach, followed by three washes with sterile ultrapure H_2_O), and stratified (dark, 4°C, 2 d). Liquid cultures for hormone measurements were grown in 6-well plates (1 mg seeds in 7 mL ½ MS medium (Murashige and Skoog 1962) (Merck, M5519) with 1% (w/v) sucrose (Merck S7903) and 0.05% (w/v) 4-morpholineethanesulfonic acid sodium salt (Merck, M3058) per well) and in 250 mL Erlenmeyer flasks for IP-LC-MS/MS (30 mg seeds in 125 mL ½ MS per flask) in 16 h light/8 h dark conditions for 7 d at 130 rpm. Seedlings used for microscopy were grown on ½ MS with 1% (w/v) agar (Merck, A1296) for 6 d, in 16 h light/8 h dark conditions.

### Plant stress treatments

*A. thaliana* stress treatments were started at the start of the photoperiod. The growth medium was exchanged for ½ MS medium (with 0.1% (v/v) DMSO) containing one of the respective treatments: 600 nM ISX (Merck, 75772), 300 mM sorbitol (D-sorbitol, Merck, S6021), 600 nM ISX and 300 mM sorbitol combined, or no additives (as control). After 4 h, the plant material was collected, briefly rinsed with ultrapure H_2_O, dried by dipping on tissue paper, and immediately flash-frozen in liquid nitrogen. Samples were stored at -80°C until further use.

### Cloning and creation of stable *A. thaliana* lines

The plasmid kit used for generation of plant transformation constructs was a gift from Jan Lohmann (Addgene kit #1000000036) (Lampropoulos et al. 2013). The promotor *THE1_pro_* was cloned into pGGA000. The open reading frame (ORF) of *THE1*_Δ*kinase*_ was amplified from *the1-4* genomic DNA and cloned into pIB/V5-His-TOPO™ with the pIB/V5-His TOPO TA Expression Kit (Thermo Fisher Scientific). The *Bsa*I restriction enzyme recognition site was removed by changing nucleotide 606 from T to A using the Q5 Site-Directed Mutagenesis Kit (New England Biolabs) and the ORF of *THE1*_Δ*kinase*_ was cloned into pGGC000. An 8-alanine-linker (*8A-linker*) was added to *eGFP* (amplified from pGGD001) and cloned into pGGD000. Plasmids A-F (using pGGB003, pGGE009, pGGF007) were assembled with pGGZ003. *THE1*_Δ*ECD*_ was obtained by gene synthesis, combining *THE1* nucleotides 1-72 (coding for the signal peptide, predicted with SignalP 6.0 (Teufel et al. 2022)), nucleotides 1162-2565 (including the transmembrane domain, predicted with TMHMM 2.0 (Möller et al. 2001)), and the *8A-linker-eGFP* (*8A-GFP*) in the assembled plasmid. All constructs were verified by sequencing.

Attempts to clone the full length *THE1* and *THE1_G37D_*ORF into pGGC000 resulted either in no colony growth after transformation of *E. coli* or mutations causing amino acid changes. Hence, THE1 and *THE1_G37D_* were cloned in two parts: *THE1.pt1* (amplified from pGGZ003 containing *THE1_pro_:THE1*_Δ*ECD*_*-GFP)* or *THE1_G37D_.pt1* (amplified from *the1-1* gDNA and *Bsa*I site removal with Q5 Site-Directed Mutagenesis Kit), corresponding to nucleotides 1-1647, in pGGB000, and *THE1.pt2,* corresponding to nucleotides 1644-2565, in pGGC000. The overlap of nucleotides 1644-1647 corresponds to the overhang GGCT of the GreenGate cloning system between plasmids B and C, allowing for seamless fusion and reconstitution of the *THE1* or *THE1_G37D_* sequence during the assembly of plasmids A-F with pGGZ003. The assembled plasmids were purified with the DNA Clean & Concentrator-5 kit (Zymo Research; elution with ultrapure H_2_O) and were directly transformed into electrocompetent *A. tumefaciens* GV3031. *THE1* construct integrity was confirmed by sequencing plasmids produced in *E. coli* or, in the case of full-length *THE1* and *THE1_G37D_*, amplified from individual *A. tumefaciens* colonies (Figs. S11-13).

For stable expression in *A. thaliana*, *the1-6* plants (loss of function, S53STOP (Merz et al. 2017)) were transformed with *A. tumefaciens* by floral dip (Clough and Bent 1998) and selected by kanamycin resistance (Harrison et al. 2006) and presence of GFP signal to homozygosity. We obtained two independent lines for THE1-GFP and THE1_K636E_-GFP, three independent lines for THE1_Δkinase_-GFP, and one independent line for THE1_G37D_-GFP and THE1_ΔECD_-GFP. *THE1* and *THE1_G37D_* construct integrity *in planta* was monitored by sequencing PCR products of *THE1* and *THE1_G37D_* amplified from *A. thaliana* leaf DNA (Fig. S14).

For the kinase activity assay, the complete intracellular region (nucleotides 1321-2565) of *THE1_K636E_* and *THE1_D634N_* (amino acid exchange by Q5 Site-Directed Mutagenesis Kit in pGGZ containing *THE1_pro_:THE1*_Δ*ECD*_*-GFP)* were cloned into pGEX-6P-1. All primers used in this study are available in Table S1.

### Kinase activity assay

GST-CRK2 (pOPINK-CRK2cyto), GST-CRK2_DN_ (pOPINK-CRK2cytoKD, D450N) (Kimura et al. 2020), GST-THE1_D634N_ (pGEX-6P-1-THE1_D634N_), and GST-THE1_K636E_ (pGEX-6P-1-THE1_K363E_) were expressed in *E. coli* expression strain SHuffle T7 (New England Biolabs, C3026J). Bacterial culture was grown to a final OD_600_ of 0.4-0.6, induced with 400 µM isopropyl β-D-1-thiogalactopyranoside and protein expressed over night at 18°C, 160 rpm. Recombinant proteins were purified with glutathione sepharose 4B beads (GE Healthcare, 17-0756-01) and eluted (20 mM reduced glutathione, 50 mM Tris/HCl pH 8.0, 200 mM NaCl and 0.1% Triton X-100). Quality and relative quantity was evaluated based on Coomassie brilliant blue staining of proteins separated in polyacrylamide gels. The kinase assay was performed as described previously (Zeiner et al. 2026) using myelin basic protein (Sigma-Aldrich, M1891) as substrate.

### Microscopy

Microscopy images were taken with a Zeiss LSM800 in airyscan mode (Fig. 1; Figs. S5 and S6) or Zeiss Axio Observer 7 (Fig. S9). Propidium iodide (PI) (Sigma-Aldrich, P4170) staining in Fig. 1c was performed with 15 µM PI in ½ MS for 15 min. Plants in Fig. 1d were stained as in Fig. 1d with the addition of 450 mM sorbitol to induce plasmolysis. GFP was excited at 488 nm with laser power not exceeding 3% and emission measured at 500-535 nm. Bleed-through signal from chloroplast autofluorescence was imaged by 488 nm excitation and 640-700 nm emission (Fig. S5). PI was excited at 561 nm with laser power not exceeding 2.5% and emission 590-650 nm. Objectives used: EC Plan-Neofluar 2.5x/0.085 M27 (whole seedling Fig. 1b, Fig. S5a), C-Apochromat 10x/0.45 W (Root tip, elongation zone, and hypocotyl Fig.1c, Fig. S5b), C-Apochromat 40x/1.20 W (epidermis Fig. 1b; Fig. 1c), Plan-Apochromat 63x/1.40 Oil (stomata Fig. 1b; Fig. 1d). Settings for Fig. S9 were excitation: 493, emission: 492-678, objective: Plan apochromat 40X/0.95 Korr M27, and Fig. S9 images are a maximum projection of 10 slices (0.3 µm depth/slice). Images were analyzed with Zen Blue V3.12 software (Carl Zeiss) and ImageJ V1.54p (Schneider et al. 2012).

**Figure 1.**
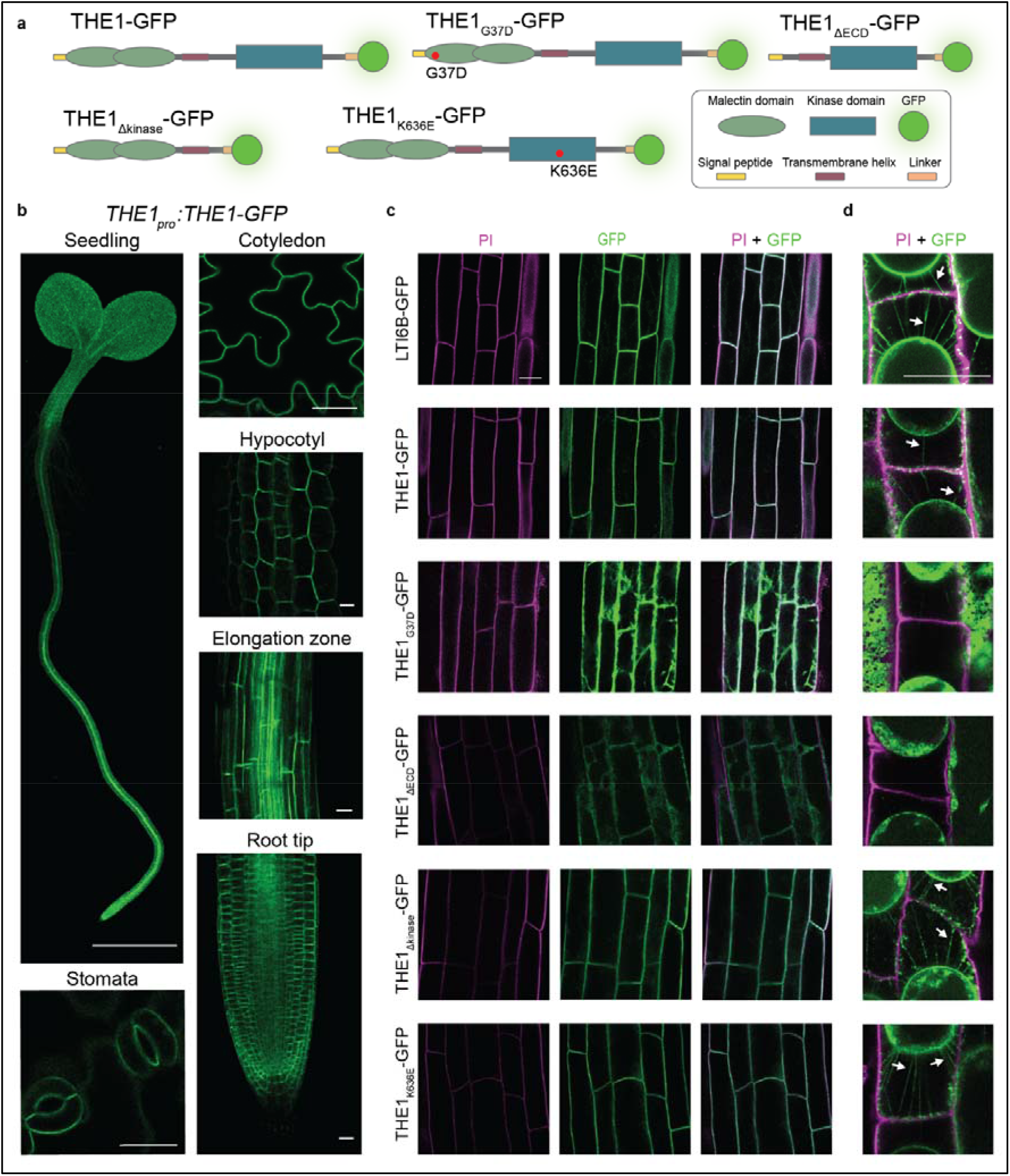
Overview of the localization of different THE1-GFP fusion constructs in six-day-old *A.thaliana* seedlings. **a)** Schematic representation of the different THE1-GFP variants. **b)** Confocal microscopy-derived images of THE1-GFP expressed under control of the native *THE1* promoter in whole seedlings, abaxial cotyledon epidermis, root tip, root elongation zone, etiolated hypocotyl, and stomata. **c)** Localization of the plasma membrane marker LTI6B-GFP and the different THE1-GFP variants in cortex/epidermal cells of the root elongation zone. **d)** LTI6B-GFP and THE1-GFP variants in plasmolyzed epidermal cells in the root elongation zone. Arrows highlight exemplary Hechtian strands. Scale bar seedling (b): 1 mm, other scalebars: 20 µm.

### JA quantification

Phytohormone extraction and quantification was based on Forcat et al. (2008). Harvested and frozen plant material was freeze-dried for 24 h. In a TissueLyzer (Qiagen, PTFE blocks pre-cooled at -80°C), 6.5 ± 0.2 mg of freeze-dried sample with one stainless steel bead (Ø 5 mm, Qiagen, #69989) was homogenized (25 Hz, 2 min), and centrifuged (17000 x g, 4°C, 2 min). 400 µL of extraction buffer (10% (v/v) methanol, 1% (v/v) acetic acid, 50 ng/mL d5-JA (CDN Isotopes)) was added, resuspended by vortexing and mixed in the TissueLyzer (25 Hz, 2 min). After incubation on ice for 30 min, debris was pelleted by centrifugation (17000 x g, 4°C, 10 min) and the supernatant was cleared through PTFE syringe filters (VWR, Ø 13 mm, 0.2 µm pore size, #514-1275).

120 µL sample was used for ultra-performance liquid chromatography-tandem mass spectrometry (UPLC-MS/MS). The analyses were performed on an Acquity I-Class UPLC system coupled to a Xevo TQ-XS triple quadrupole mass spectrometer (both from Waters Corporation), operating in negative electrospray ionization mode with multiple reaction monitoring, as described by Bacete et al. (2022) with minor modifications. Chromatographic separation was achieved on an ACQUITY UPLC BEH C18 column (1.7 μm, 2.1 × 100 mm). The mobile phase consisted of water (A) and acetonitrile (B), both containing 0.1% formic acid, at a flow rate of 0.4 mL/min. The column temperature was maintained at 30°C, and the injection volume was 5 μL. The linear gradient was set as follows: 0-1.2 min, 20% B; 1.2-4.0 min, linear ramp to 95% B; 4.0-5.0 min, hold at 95% B; 5.0-6.0 min, return to 20% B; and 6.0-7.0 min, re-equilibration at 20% B. Quantification of JA was performed using isotopic dilution calibration curves ranging from 0.2 to 1000 ng/mL. The limit of quantification was 0.2 ng/mL.

### Immunoprecipitation and proteomic analysis by mass spectrometry

Frozen samples were pulverized in liquid nitrogen with mortar and pestle. All following procedures were conducted in the cold room at 4°C. In 50 ml reaction tubes, 10 g of plant powder was combined with 20 mL of pre-cooled lysis buffer (50 mM Tris-HCl pH 7.5, 150 mM NaCl, 10 mM MgCl_2_, 5 mM EDTA, 10% (v/v) glycerol, 1% (v/v) IGEPAL CA-630, 2 mM dithiotreitol (DTT), 333 µl protease inhibitor cocktail (Sigma-Aldrich, P9599), and 1x PhosSTOP (Roche, 4906837001)). Samples were resuspended by vortexing and mixed for 1 h on a roller shaker. Insoluble fractions were pelleted by centrifugation (4000 x g, 4°C, 15 min) and filtered through 22-25 µm pore size Miracloth (Calbiochem, 475855). Samples were incubated overnight with 50 µL of ChromoTek GFP-Trap magnetic agarose beads (Proteintech, gtma). Using a magnetic rack, the GFP-Trap beads were washed twice with 1 mL lysis buffer and thrice with washing buffer (50mM Tris-HCl pH 7.5, 150 mM NaCl). Lastly, the GFP-Trap beads were resuspended in 40 µL 50 mM Tris-HCl and stored at -20°C until proteomics analysis.

MS analysis was performed by the Proteomics and Modomics Experimental Core Facility of the Norwegian University of Science and Technology. A 30 µl sample was combined with 150 µL 0.1 M ammonium bicarbonate. Proteins were reduced with 1.5 µl 1M DTT (55°C, 30 min), alkylated with 15 µl 200 mM iodoacetamide (30 min, 20°C, dark), and digested with 500 ng trypsin (Thermo Fisher Scientific, 90057) (37°C, overnight). Peptides were desalted using C18 spin columns (Thermo Fisher Scientific, 89870), dried in a SpeedVac (Thermo Fisher Scientific) and resuspended in 0.1% formic acid before performing LC-MS/MS (timsTOF Pro 2, DIA PASEF mode) with the nano-elute liquid chromatography system (Bruker Daltonics). The peptides were separated for 75 min using a 150 µm x 25 cm x 1.9 µm PepSep 25 (Bruker Daltonics) with running buffers A (0.1% formic acid) and B (0.1% formic acid in acetonitrile) with a gradient of 2-40%. Source conditions: 1600 V capillary voltage, 3.0 L/min dry gas, 180°C dry temperature. MS1 and MS2 spectra m/z range: 100-1700, collision energy linearly interpolated between 59 eV (1/K₀ = 1.60 V·s/cm²) and 20 eV (1/K₀ = 0.6 V·s/cm²).

Raw MS data were processed using DIA-NN v2.2.0 Academia (Demichev et al. 2020). DiaPASEF runs were analyzed using a predicted spectral library reannotated against the UniProt *A. thaliana* reference proteome (UP000006548, downloaded April 2026) supplemented with contaminant protein sequences. Settings were: trypsin specificity: ≤2 missed cleavages, peptide lengths: 7-30 amino acids, charge: 1-4, precursor m/z: 300-1800, fragment m/z: 200-1800, fixed modification: cysteine carbamidomethylation, variable modifications (max 3/peptide): methionine oxidation and protein N-terminal acetylation, mass accuracy settings: fixed at 15 ppm for both precursor and fragment ions. Peptidoform scoring, retention-time profiling, and high-accuracy QuantUMS quantification were enabled (Grossmann et al. 2026). Protein inference was performed using proteotypic peptides. False discovery rates (FDRs) were controlled at 1% at the precursor and protein levels. Match-between-runs was used to reduce missing values, transferring peptide IDs across runs after verifying chromatographic and ion-mobility evidence. Protein group intensities were normalized by DIA-NN label-free quantification (Data S2; Cox et al. 2014). The R-package DEP2 v0.5.28.2 was used to identify differentially enriched protein groups (Feng et al. 2023). Protein groups were filtered (peptide number >2, values present in at least one triplicate sample group ≥ 2), variance normalized (function normalize_vsn), imputed (function MinDet), and significance tested (*limma* (Ritchie et al. 2015) with Strimmer’s Q-value FDR correction (Strimmer 2008), fold-change >2, and FDR <0.05). Heatmaps were made with the R-package ComplexHeatmap v2.14.0 (Gu 2022). GO enrichment analysis was performed with ShinyGO v0.77 (Ge et al. 2020).

## Results

### Mutations in the THE1 ectodomain lead to mislocalization

We generated a set of *THE1-GFP* fusion constructs in which different parts of THE1 are modified to investigate the localization of wild-type and mutant THE1 versions (Fig. 1a). These constructs include *THE1-GFP* (full length *THE1*), *THE1_K636E_-GFP*, *THE1*_Δ*kinase*_*-GFP* (deleted kinase domain, corresponds to *the1-4*), *THE1_G37D_-GFP* (G37D mutation in the first malectin-like domain, corresponds to *the1-1*), and *THE1*_Δ*ECD*_*-GFP* (deleted ectodomain but signal peptide is retained). The THE1_K636E_ variant is mutated in the catalytic-loop lysine K636 of the HRD-x-K motif (Hanks et al. 1988; Kornev et al. 2006), and structural modelling predicted that this mutation impairs kinase activity (Fig. S1). However, an *in vitro* kinase assay performed on the intracellular part of THE1_K636E_ showed autophosphorylation activity (weak in comparison to the cytosolic part of CYSTEINE-RICH RECEPTOR-LIKE PROTEIN KINASE 2 (Kimura et al. 2020) used as positive control), while little to no general substrate (myelin basic protein) phosphorylation was observed (Fig. S2). Due to the detected residual kinase activity, we will hereafter refer to THE1_K636E_ as a kinase-mutated variant. All *in planta* constructs were expressed under the control of the native *THE1* promoter (*THE1_pro_*) in *the1-6* (S53STOP (Merz et al. 2017)) *A. thaliana* background. The *the1-6* background was favored over the *the1-1* background because *Cr*RLK1Ls may dimerize (Kong et al. 2023; Li et al. 2026a). In the *the1-1* background a THE1_G37D_ loss-of-function protein is present which may interact with THE1-GFP and impair its function. Only 52 N-terminal residues (of which 24 make up the secretion peptide) are produced in *the1-6* background (Merz et al. 2017), corresponding to a knock-out of *THE1*. Lines expressing *THE1-GFP*, *THE1_G37D_-GFP* or *THE1*_Δ*ECD*_*-GFP* showed no obvious phenotypic differences in four-week-old plants when compared to Columbia (Col-0), *the1-1* or *the1-6* plants (Fig. S3). We noticed that *the1-4* and lines expressing *THE1_K636E_-GFP* (two independent lines) or *THE1*_Δ*kinase*_*-GFP* (three independent lines) had consistently darker petioles than Col-0 or *the1-6* in our growth conditions (Fig. S4).

To determine the (sub)cellular localization of THE1, we performed confocal microscopy on six-day-old seedlings expressing *THE1_pro_:THE1-GFP* (Fig. 1b; Fig. S5). We observed THE1-GFP signal in both the root and the shoot. In the root, expression of THE1-GFP was most abundant in the tip and gradually reduced in the basal direction. In the shoot, expression appeared to be equally distributed across hypocotyl and cotyledon. We then focused on the cotyledon epidermis, stomata, hypocotyl epidermis and cortex, elongation zone medial plane, and root tip medial plane (Fig. 1c and d). In all tissues investigated, THE1-GFP localized to the cell periphery, suggesting an association with the plasma membrane. Next, we verified that THE1 localizes to the plasma membrane and determined how different modifications at the protein level affect THE1 localization. We used the LTI6B-GFP (Cutler et al. 2000) plasma membrane marker line to visualize the plasma membrane and establish the localization of the THE1-GFP variants (Fig. 1c). THE1-GFP, THE1_K636E_-GFP and THE1_Δkinase_-GFP had a comparable subcellular localization as LTI6B-GFP, suggesting that THE1 localizes to the plasma membrane. In contrast, THE1_G37D_-GFP and THE1_ΔECD_-GFP seemed to be intracellularly localized. To determine the localization of the THE1 variants more clearly, we induced plasmolysis and monitored the GFP signal (Fig. 1d; Fig. S6). GFP signal was clearly observed in the plasma membrane and Hechtian strands in plants expressing LTI6B-GFP, THE1-GFP, THE1_K636E_-GFP, and THE1_Δkinase_-GFP, confirming that these proteins are indeed localized to the plasma membrane. These observations also show that neither the kinase domain nor enzymatic activity of THE1 are required for correct localization. However, in plants expressing THE1_G37D_-GFP or THE1_ΔECD_-GFP the signal from Hechtian strands was either very weak or absent. These results demonstrate that G37D mutation or ectodomain removal impairs localization of THE1 to the plasma membrane.

### THE1 requires an intact ectodomain but no kinase to induce JA production

*A. thaliana* seedlings encoding the *the1-1* loss-of-function allele produce less JA after ISX treatment, whereas seedlings carrying the *the1-4* allele are known to produce more JA than Col-0 seedlings following CWI impairment by ISX (Engelsdorf et al. 2018; Bacete et al. 2022). To verify that the JA production phenotype is also present in the *the1-6* loss-of-function background and test if complementation with the *THE1-GFP* constructs rescues the phenotype we measured JA levels after 8 hours of treatment with ISX or DMSO (mock-treated control, as DMSO is used as solvent for ISX, Fig. 2a and b). JA levels are low in control conditions for most lines; *the1-4* as well as *THE1*_Δ*kinase*_*-GFP* and *THE1_K636E_-GFP* in the *the1-6* background show slightly elevated JA levels. After ISX treatment, Col-0 seedlings show a clear induction of JA synthesis. JA levels in ISX-treated *the1-1* and *the1-6* seedlings were much lower than in Col-0. The strongly reduced JA production phenotype in *the1-1* and *the1-6* indicates they are both loss-of-function alleles. JA levels in ISX-treated *the1-4* seedlings were significantly higher than in Col-0. The introduction of *THE1-GFP* in *the1-6* plants restored JA production to Col-0 levels in one out of the two complementation lines we tested (#1.10), demonstrating that the JA production can be rescued by wild-type *THE1*. This *THE1-GFP* line was used for all further experiments. Complementation with *THE1*_Δ*kinase*_*-GFP* and *THE1_K636E_-GFP* similarly restored ISX-induced JA production, suggesting that an intact THE1 kinase domain is not required for induction of JA. Interestingly, seedlings containing the kinase-mutated *THE1_K636E_-GFP* transgene exhibited elevated JA levels (similar to *the1-4*) in control conditions when compared to Col-0 seedlings. Complementation of *the1-6* with *THE1_G37D_-GFP* or *THE1*_Δ*ECD*_*-GFP* did not result in a rescue of the *the1-6* phenotype, indicating that THE1 requires a functional ectodomain to induce JA production.

**Figure 2.**
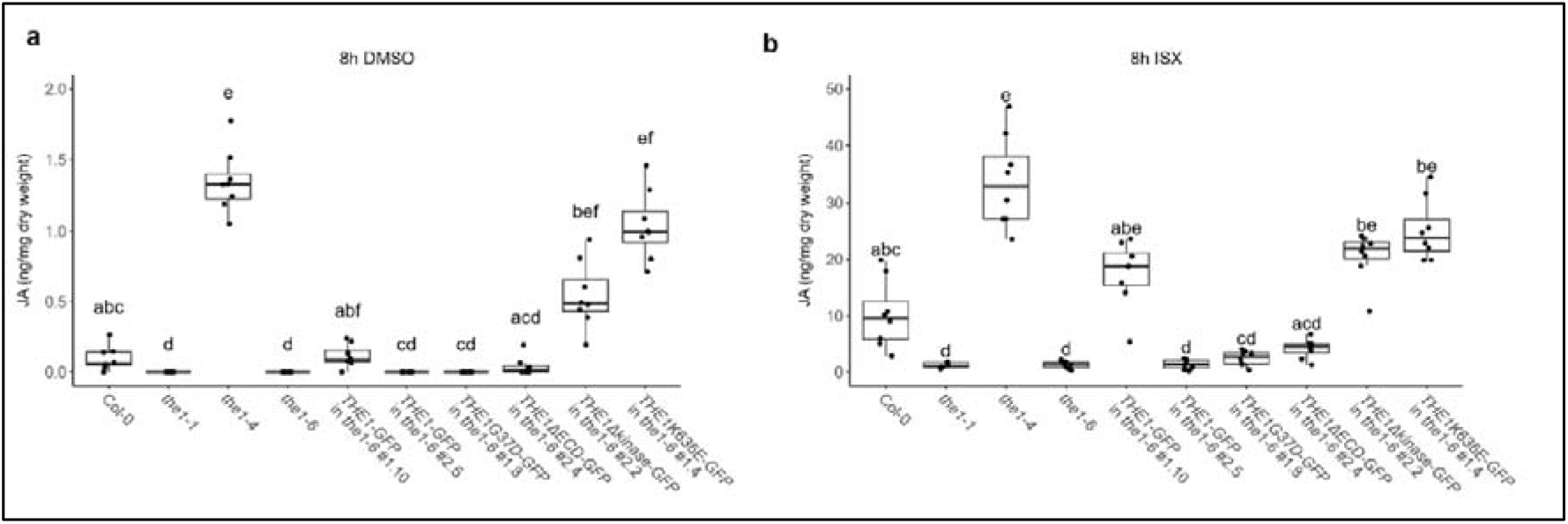
Jasmonic acid (JA) levels in Col-0, *the1* mutant lines and *THE1-GFP* transgenic *A. thaliana* in response to CWI impairment. JA levels were quantified by ultra-performance liquid chromatography-tandem mass spectrometry following 8 hours treatment of seven-day-old seedlings with **a)** DMSO (mock) or **b)** ISX. Different letters indicate significance as determined by Kruskal-Wallis test followed by Dunn’s post-hoc test (p < 0.05). Boxplots indicate the median (center line), Q1-Q3 interquartile range (box limits), and 1.5x interquartile range (whiskers). Note that the Y-axis scale differs between (a) and (b).

### THE1 interacts with a functionally diverse set of proteins

To identify proteins involved in THE1 activation, regulation and signaling, we performed IP-LC-MS/MS on seven-day old THE1-GFP-expressing *A. thaliana* seedlings to which no stress treatment was applied, using LTI6B-GFP-expressing plants (Cutler et al. 2000) as control to identify unspecific interactors (Fig. 3a and b; Data S1). The experiment was performed twice, and peptides were measured on independent MS runs to identify the most reliable candidates. We identified 138 putative interactors: 82 in experiment 1 (Fig. 3a), 97 in experiment 2 (Fig. 3b) and 41 identified in both experiments (Fig. 3c; Table 1).

**Figure 3.**
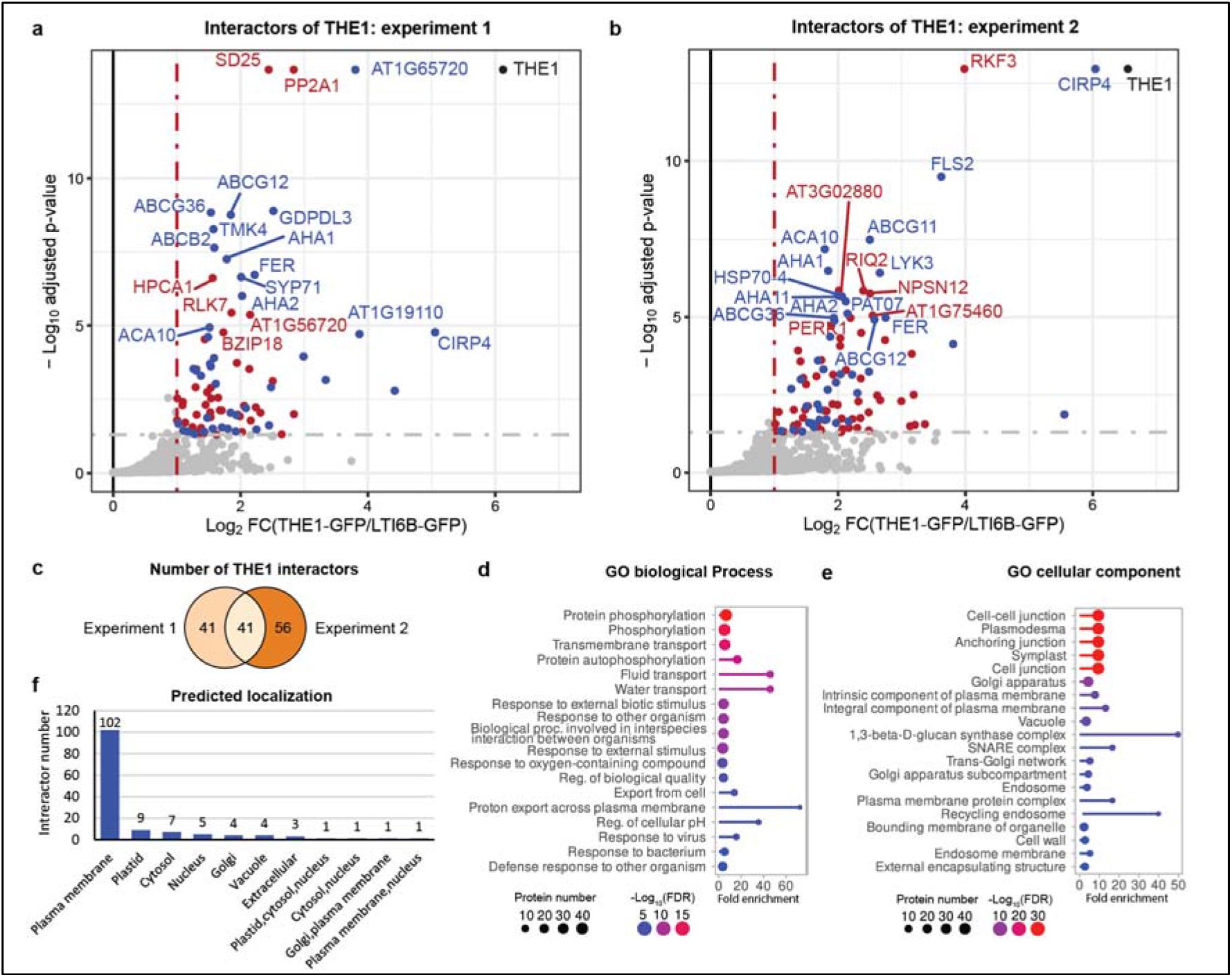
Putative interactors of THE1 identified by immunoprecipitation-liquid chromatography-tandem mass spectrometry (IP-LC-MS/MS) using THE1-GFP *A. thaliana* seedlings in non-stress conditions. **a)** and **b)** Putative interactors of THE1 as determined by two independent IP-LC-MS/MS experiments (n=3 per experiment). Putative interactors were identified by enrichment analysis with the R-package *limma* (Ritchie et al. 2015) and false discovery rate (FDR) correction using the Strimmer’s Q-value method. Proteins with a fold-change (FC) >2 and FDR-adjusted P-value <0.05 were considered significantly enriched. Blue dots: interactors that were significantly enriched in both experiments; red dots: significantly enriched interactors identified in one of two experiments; grey dots: non-significant enriched proteins; black dot: THE1 bait protein. Names of the 20 most significant interactors are indicated. **c)** Venn diagram showing the number and overlap of significant interactors in the two experiments. **d)** and **e)** GO term enrichment analysis showing the biological processes (d) and cellular component (e) associated with the 138 unique interactors pooled from both experiments. Circle size: number of interactors associated with the GO term; circle color: -log_10_ of FDR-adjusted p-values.

**Table 1.** Recurring putative interactors of THE1 identified by immunoprecipitation. -**liquid chromatography**-tandem mass spectrometry (IP-LC-MS/MS) using THE1-GFP *A. thaliana* seedlings in non-stress conditions. The table shows putative THE1 interactors detected in two independent IP-LC-MS/MS experiments (n = 3 per experiment). Columns show the TAIR identifier (ID), UniProt ID, protein symbol, full protein name, (predicted) protein function according to UniProt, (predicted) protein localization according to the subcellular localization database for *A. thaliana* proteins (SUBA5) (Hooper et al. 2017), total and unique peptides detected for the protein in experiment (Exp.) 1 and 2, and log_2_ of the fold change (FC), where FC = (protein group intensity in THE1-GFP/protein group intensity in LTI6B-GFP). *: SHOU4 was detected as a protein group together with its homolog SHOU4-LIKE in Exp.1.

| TAIR ID | UniProt iD | Protein symbol | Full protein name | (Predicted) protein function | (Predicted) localization | Peptides (Unique) Exp. 1 | Peptides (Unique) Exp. 2 | Log2(FC) Exp.1 | Log2(FC) Exp.2 |
|---|---|---|---|---|---|---|---|---|---|
| AT2G39360 | O80623 | CVY1 | CURVY1 | *Cr*RLK1L receptor kinase | Plasma membrane | 8(8) | 12(12) | 1,80 | 1,51 |
| AT3G51550 | Q9SCZ4 | FER | FERONIA | *Cr*RLK1L receptor kinase | Plasma membrane | 21(18) | 25(22) | 2,22 | 2,75 |
| AT1G28440 | Q9SGP2 | HSL1 | HAESA-LIKE 1 | Receptor kinase | Plasma membrane | 6(5) | 8(7) | 1,43 | 2,30 |
| AT1G51940 | F4IB81 | LYK3 | LYSM DOMAIN RECEPTOR-LIKE KINASE 3 | Receptor kinase | Plasma membrane | 3(3) | 3(3) | 4,42 | 2,66 |
| AT1G66150 | P43298 | TMK1 | TRANSMEMBRANE KINASE 1 | Receptor kinase | Plasma membrane | 5(4) | 9(8) | 2,25 | 2,16 |
| AT3G14350 | Q9LUL4 | SRF7 | Protein STRUBBELIG-RECEPTOR FAMILY 7 | Receptor kinase | Plasma membrane | 3(3) | 9(9) | 2,45 | 1,71 |
| AT3G14840 | C0LGN2 | LIK1 | LYSM RLK1-INTERACTING KINASE 1 | Receptor kinase | Plasma membrane | 26(23) | 29(26) | 1,31 | 1,79 |
| AT3G23750 | Q9LK43 | TMK4 | TRANSMEMBRANE KINASE 4 | Receptor kinase | Plasma membrane | 14(14) | 23(22) | 1,57 | 1,88 |
| AT4G23180 | Q8GYA4 | CRK10 | CYSTEINE-RICH RECEPTOR-LIKE PROTEIN KINASE 10 | Receptor kinase | Plasma membrane | 8(5) | 15(10) | 2,48 | 1,62 |
| AT4G39400 | O22476 | BRI1 | BRASSINOSTEROID INSENSITIVE 1 | Receptor kinase | Plasma membrane | 7(6) | 8(7) | 1,94 | 1,54 |
| AT5G16590 | Q9FMD7 | AT5G16590 | AT5G16590 | Receptor kinase | Plasma membrane | 9(9) | 25(18) | 1,56 | 1,77 |
| AT5G46330 | Q9FL28 | FLS2 | FLAGELLIN SENSITIVE 2 | Receptor kinase | Plasma membrane | 9(9) | 10(10) | 2,99 | 3,62 |
| AT1G17840 | Q8RXN0 | ABCG11 | ABC TRANSPORTER G FAMILY MEMBER 11 | ABC transporter | Plasma membrane | 22(21) | 27(26) | 1,58 | 2,50 |
| AT1G51500 | Q9C8K2 | ABCG12 | ABC TRANSPORTER G FAMILY MEMBER 12 | ABC transporter | Plasma membrane | 12(9) | 23(19) | 1,85 | 2,57 |
| AT1G59870 | Q9XIE2 | ABCG36 | ABC TRANSPORTER G FAMILY MEMBER 36 | ABC transporter | Plasma membrane | 42(18) | 61(30) | 1,53 | 1,93 |
| AT2G26910 | O81016 | ABCG32 | ABC TRANSPORTER G FAMILY MEMBER 32 | ABC transporter | Plasma membrane | 23(22) | 36(34) | 1,61 | 1,94 |
| AT3G28860 | Q9LJX0 | ABCB19 | ABC TRANSPORTER B FAMILY MEMBER 19 | ABC transporter | Plasma membrane | 26(23) | 33(31) | 1,49 | 1,97 |
| AT3G62150 | Q9M1Q9 | ABCB21 | ABC TRANSPORTER B FAMILY MEMBER 21 | ABC transporter | Plasma membrane | 18(8) | 28(16) | 1,53 | 1,11 |
| AT4G25960 | Q8LPK2 | ABCB2 | ABC TRANSPORTER B FAMILY MEMBER 2 | ABC transporter | Plasma membrane | 23(20) | 35(29) | 1,59 | 2,22 |
| AT2G18960 | P20649 | AHA1 | AUTOINHIBITED H+-ATPase 1 | H<sup>+</sup>-ATPase | Plasma membrane | 42(4) | 36(5) | 1,78 | 1,85 |
| AT4G30190 | P19456 | AHA2 | AUTOINHIBITED H+-ATPase 2 | H<sup>+</sup>-ATPase | Plasma membrane | 19(0) | 17(0) | 2,02 | 2,15 |
| AT5G57350 | P20431 | AHA3 | AUTOINHIBITED H+-ATPase 3 | H<sup>+</sup>-ATPase | Plasma membrane | 7(2) | 24(6) | 1,18 | 1,68 |
| AT5G62670 | Q9LV11 | AHA11 | AUTOINHIBITED H+-ATPase 11 | H<sup>+</sup>-ATPase | Plasma membrane | 32(14) | 42(18) | 1,52 | 2,06 |
| AT3G09740 | Q9SF29 | SYP71 | SYNTAXIN-71 | Syntaxin | Plasma membrane | 8(8) | 15(15) | 2,01 | 1,84 |
| AT3G11820 | Q9ZSD4 | SYP121 | SYNTAXIN-121 | Syntaxin | Plasma membrane | 7(7) | 10(10) | 2,08 | 1,82 |
| AT5G08080 | Q8VZU2 | SYP132 | SYNTAXIN-132 | Syntaxin | Plasma membrane | 11(11) | 14(13) | 3,34 | 2,04 |
| AT5G53480 | Q9FJD4 | KPNB1 | KARYOPHERIN-β 1 | Nuclear transport receptor protein | Plastid,cytosol,nucleus | 17(17) | 17(17) | 1,26 | 1,41 |
| AT3G08943 | F4IYK3 | AT3G08943 | AT3G08943 | Nuclear transport receptor protein | Cytosol,nucleus | 8(3) | 10(5) | 1,48 | 1,30 |
| AT5G57110 | Q9LF79 | ACA8 | AUTOINHIBITED CA<sup>2+</sup> -ATPASE ISOFORM 8 | Ca<sup>2+</sup> efflux pump | Plasma membrane | 16(15) | 28(25) | 1,38 | 1,69 |
| AT4G29900 | Q9SZR1 | ACA10 | AUTOINHIBITED CA<sup>2+</sup> -ATPASE ISOFORM 10 | Ca<sup>2+</sup> efflux pump | Plasma membrane | 18(17) | 23(23) | 1,51 | 1,79 |
| AT1G30360 | Q9C8G5 | OSCA3.1 | HYPEROSMOLALITY-GATED CA<sup>2+</sup>-PERMEABLE CHANNEL 3.1 | Ion channel | Plasma membrane | 17(17) | 24(24) | 1,10 | 1,44 |
| AT4G35100 | P93004 | PIP2-7 | PLASMA MEMBRANE INTRINSIC PROTEIN 2-7 | Aquaporin | Plasma membrane | 3(3) | 3(3) | 1,84 | 1,97 |
| AT4G03550 | Q9ZT82 | CALS12 | CALLOSE SYNTHASE 12 | Callose synthase | Plasma membrane | 24(21) | 32(27) | 1,02 | 1,27 |
| AT5G60710 | Q9FF49 | CIRP4 | CATALASE2 INTERACTING RING PROTEIN 4 | Probable ubiquitin ligase | Plasma membrane | 14(14) | 24(24) | 5,06 | 6,04 |
| AT1G19110 | Q8L798 | AT1G19110 | INTER-ALPHA-TRYPSIN INHIBITOR HEAVY CHAIN-LIKE PROTEIN | Probable ubiquitin ligase | Plasma membrane | 13(13) | 19(19) | 3,87 | 5,55 |
| AT4G30340 | F4JQ95 | DGK7 | DIACYLGLYCEROL KINASE 7 | Diacylglycerol kinase | Cytosol | 8(8) | 11(10) | 1,71 | 1,60 |
| AT3G12580 | Q9LHA8 | HSP70-4 | HEAT SHOCK 70 KDA PROTEIN 4 | Heat shock protein | Cytosol | 18(11) | 26(16) | 1,27 | 1,99 |
| AT3G26935 | Q0WQK2 | PAT07 | PROTEIN S-ACYLTRANSFERASE 7 | Probable protein S-acyltransferase | Plasma membrane | 4(4) | 8(8) | 1,30 | 2,12 |
| AT4G26690 | Q9SZ11 | GDPDL3 | GLYCEROPHOSPHORYL DIESTER PHOSPHODIESTERASE-LIKE PROTEIN 3 | Glycerophosphodiester phosphodiesterase | Plasma membrane | 11(10) | 9(8) | 2,52 | 2,49 |
| AT1G65720 | Q9SHY3 | AT1G65720 | AT1G65720 | Unknown | Plasma membrane | 5(5) | 6(6) | 3,80 | 3,81 |
| AT1G78880 | Q9ZVA7 | SHOU4 | SHOU4 | Unknown | Plasma membrane | 6(0)* | 12(4) | 1,92 | 1,72 |

The 41 recurring THE1 interactors include proteins of known and unknown functions (Table 1). We observed interactors belonging to the *Cr*RLK1L family, such as FER (Huck et al. 2003; Escobar-Restrepo et al. 2007) and CURVY1 (Gachomo et al. 2014), and interactors from other RK families such as FLAGELLIN SENSITIVE 2 (FLS2) (Gómez-Gómez and Boller 2000) and BRI1 (Li and Chory 1997). Some interactors have functions in transmembrane transport, such as the calcium pumps AUTOINHIBITED CA^2+^-ATPASE (ACA) 8 and -10 (Frei dit Frey et al. 2012), the mechanosensitive ion channel REDUCED HYPEROSMOLALITY-INDUCED Ca^2+^ INCREASE 3.1 (OSCA3.1) (Zhang et al. 2018b), and the aquaporin PLASMA MEMBRANE INTRINSIC PROTEIN 2-7 (PIP2-7) (Hachez et al. 2014). Seven ATP-binding cassette (ABC) transporters were observed: ABC TRANSPORTER FAMILY B (ABCB) 2, -19, and -21 (Devi et al. 2025), and ABC TRANSPORTER FAMILY G (ABCG) 19, -21 -32, and -36 (Gräfe and Schmitt 2021; Lou et al. 2026). Four different AUTOINHIBITED H^+^-ATPases (AHAs) were observed: AHA1, -2, -3, and -11 (Li et al. 2022). Two recurrent interactors have been associated with in regulation of cellulose biosynthesis: SHOU4 (Polko et al. 2018) and GLYCEROPHOSPHORYL DIESTER PHOSPHODIESTERASE-LIKE PROTEIN 3 (GDPDL3), also known as SHAVEN3 (Hayashi et al. 2008). The protein CATALASE 2 INTERACTING RING PROTEIN 4 (CIRP4) was detected twice as one of the most significant and most enriched interactors. CIRP4 is a homolog of the E3 ubiquitin ligase CIRP1 (54% shared sequence identity) (Yang et al. 2025). Other putative interactors of interest are DIACYLGLYCEROL KINASE 7 (DGK7) (Gómez-Merino et al. 2004), which converts diacylglycerol into the second messenger phosphatidic acid (PA) (Gómez-Merino et al. 2005), and KARYOPHERIN-β 1 (KPNB1) and its homolog AT3G08943 (59.9% shared amino acid sequence identity) that are involved in nuclear import (Oh et al. 2020). The interactor AT1G65720 is strongly enriched but has no described function. An InterPro database (Blum et al. 2026) query showed that AT1G65720 has a predicted signal peptide for secretion (residue 1-25), an extracellular region with no homology to known domains (residue 26-106), a transmembrane helix (residue 107-135), and an intracellular region (residue 136-180) (Fig. S7). Several of these interactors may be involved in known roles of THE1 in stress signaling and determining cell wall structure, while others associate THE1 with pathways previously not described for THE1 such as ubiquitin signaling, PA signaling, and nuclear import.

Next, we performed a Gene Ontology (GO) term enrichment analysis to predict the biological process and cellular components associated with the 138 interactors. The most significant biological process GO terms associated with the interactors include protein phosphorylation, transmembrane transport, fluid transport, and response to external biotic stimulus (Fig. 3d). These functions align with a kinase involved in biotic and abiotic stress signaling. The most significant cellular component GO terms are “Cell-cell junction”, “Plasmodesma”, “Cell junction”, “Symplast”, and “Anchoring junction” (Fig. 3e), showing that a significantly enriched fraction of THE1 interactors is associated with peripheral cellular components. This was further confirmed by using the subcellular localization database for *A. thaliana* proteins SUBA5 (Hooper et al. 2017), which showed that the majority (102 out of 138) of the interactors has a predicted plasma membrane localization. Nine interactors were predicted to localize in plastids, seven in the cytosol, five in the nucleus, four in the Golgi, four in the vacuole, and four had multiple predicted localization sites. Only three proteins had predicted extracellular localization. Thus, we mainly identified plasma membrane-localized interactors.

### THE1 mutations strongly alter the THE1 interactome

Mutations can affect protein function in different ways, including by altering interactions. In the case of THE1, a G37D mutation (*the1-1*) leads to reduced responses to cellulose biosynthesis inhibition, whereas a truncated THE1 without the kinase domain (*the1-4*) displays increased responses (Merz et al. 2017; Bacete et al. 2022). However, it is not known how these mutations affect THE1 interactions and if the increased responses in *the1-4* are due to absence of the kinase or its activity (Merz et al. 2017). We aimed to characterize the effects of G37D mutation, extracellular domain deletion, kinase deletion, and mutation of the kinase catalytic loop on the interactome of THE1. Therefore, we performed IP-LC-MS/MS using GFP-tagged THE1_G37D_, THE1_ΔECD_, THE1_Δkinase_, and THE1_K636E_ under non-stressed conditions to identify their interactomes (Fig. 4a; Data S1) and compared them to the 82 THE1 interactors identified in experiment 1 (corresponding to Fig. 3a). We decided to compare interactomes only to those THE1 interactors because all samples for the THE1 variants (except THE1_ΔECD_) were processed in the same MS batch, reducing batch effects. Out of 82 interactors, THE1 shared 18 interactors with THE1_G37D_, none with THE1_ΔECD_, six with THE1_Δkinase_, and 29 with THE1_K636E_. The majority of interactors is specific for the interaction with THE1_G37D_, THE1_ΔECD_, and THE1_Δkinase_, showing that these mutations strongly alter the THE1 interactome (Fig. 4b).

**Figure 4.**
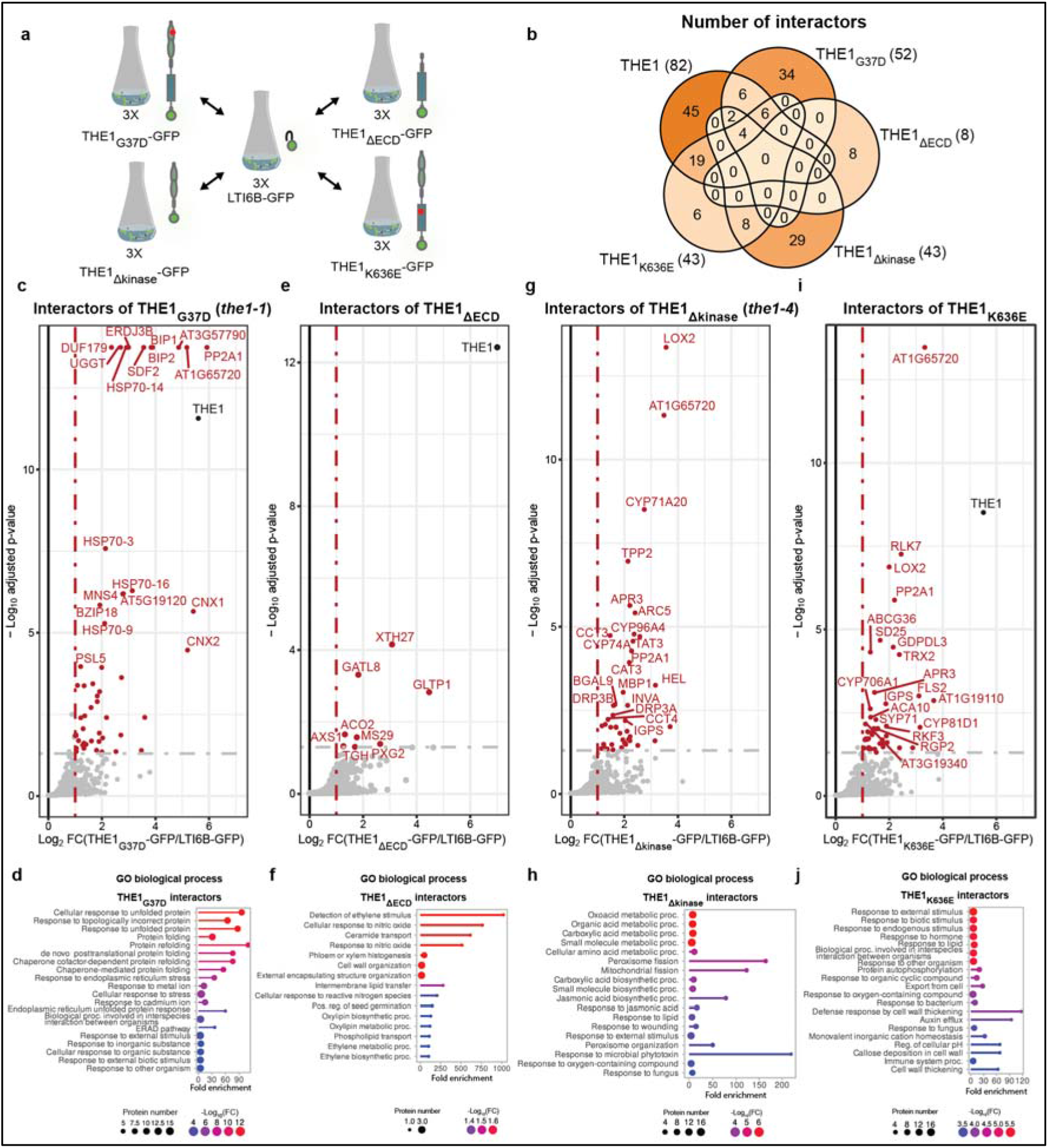
Putative interactors of THE1 variants identified by immunoprecipitation-liquid chromatography-tandem mass spectrometry (IP-LC-MS/MS) using *A. thaliana* seedlings in non-stress conditions. **a)** THE1-GFP variants or LTI6B-GFP control were used as input for IP-LC-MS/MS (n=3). **b)** Venn diagram showing number and overlap of interactors in wild-type THE1 (corresponding to Fig. 3a) and THE1 variants. The total interactor number per THE1 variant is indicated in parentheses. Interactors and interactor biological process Gene Ontology (GO) enrichment analysis of **c-d)** THE1_G37D_, **e-f)** THE1_ΔECD_, **g-h)** THE1_Δkinase_, and **i-j)** THE1_K636E_. The significance of putative interactors was determined by enrichment analysis with the R-package *limma* (Ritchie et al. 2015) and false discovery rate (FDR) correction using the Strimmer’s Q-value method. Proteins with a fold-change (FC) >2 and adjusted p-value <0.05 were considered significantly enriched. Names of the 20 most significant interactors are indicated in (c), (e), (g) and (i). FDR cutoff for pathway GO enrichment analysis: p-value <0.05. Red dots: proteins identified as significantly enriched interactors; grey dots: non-significant enriched proteins; black dot: THE1 bait protein. Circle sizes in the GO enrichment plots indicate the number of interactors associated with the GO term. Circle color: -log_10_ of FDR-adjusted p-values.

For THE1_G37D_, we detected 52 putative interactors (Fig. 3c; Data S1). The most significant THE1_G37D_ interactors include BINDING IMMUNOGLOBULIN PROTEIN (BIP) 1 and -2 (Koizumi 1996; Martínez and Chrispeels 2003), UDP-GLUCOSE:GLYCOPROTEIN GLUCOSYLTRANSFERASE (UGGT) (Blanco-Herrera et al. 2015), CALNEXIN1 (CNX) 1 and -2 (Liu et al. 2017), and ER DNAJ DOMAIN-CONTAINING PROTEIN 3B (ERDJ3B) (Shen and Hendershot 2005; Fig. 4c). The latter form core components of the ER-localized protein quality control mechanism involved in protein folding and processing of misfolded proteins (Strasser 2018). A GO analysis showed that the top biological process linked to THE1_G37D_ interactors is “Cellular response to unfolded proteins” (Fig. 4d). The top cellular component GO term associated with THE1_G37D_ interactors is “Endoplasmic reticulum lumen” (Fig. S8). These results suggest that THE1_G37D_ does not fold correctly and is retained in the ER.

For THE1_ΔECD_, eight interactors were significantly enriched (Fig. 4e; Data S1: XYLOGLUCAN ENDOTRANSGLUCOSYLASE/HYDROLASE 27 (XTH27) (Matsui et al. 2005), PEROXYGENASE 2 (PXG2) (Miklaszewska et al. 2023), GALACTURONOSYLTRANSFERASE-LIKE 8 (GATL8) (Sterling et al. 2006), GLYCOLIPID TRANSFER PROTEIN 1 (GLTP1) (West et al. 2008), 1-AMINOCYCLOPROPANE-1-CARBOXYLATE OXIDASE 2 (ACO2) (Houben and Van de Poel 2019), MITOCHONDRIAL SMALL SUBUNIT 29 (MS29) (Scarpin et al. 2023), UDP-D-APIOSE/UDP-D-XYLOSE SYNTHASE 1 (AXS1) (Mølhøj et al. 2003), and G-patch domain-containing protein TOUGH (TGH) (Calderon-Villalobos et al. 2005). While the biological process GO terms “Cell wall organization” and “External encapsulating structure organization” were enriched, this must be viewed with caution since the overall number of interactors analyzed is small. The THE_ΔECD_ interactors were not enriched for a particular cellular component GO term.

For THE1_Δkinase_, we identified 43 putative interactors (Fig. 4h; Data S1). The most significant interactors for THE1_Δkinase_ include LIPOXYGENASE 2 (LOX2) (Bell et al. 1995), cytochrome P450 family members CYP96A4 (Huang et al. 2024) and CYP74A (Laudert et al. 1996), ACCUMULATION AND REPLICATION OF CHLOROPLAST 5 (ARC5) (Gao et al. 2003), TYROSINE AMINOTRANSFERASE 3 (TAT3) (Riewe et al. 2012), MONODEHYDROASCORBATE REDUCTASE 3 (MDAR3) (Lisenbee et al. 2005), and DYNAMIN-RELATED PROTEIN 3A (DRP3A) and its homolog DRP3B (Fujimoto et al. 2009). The most significant biological process GO terms enriched amongst the THE1_Δkinase_ interactors are “Carboxylic acid metabolic process”, “Small molecule metabolic process”, “Carboxylic acid biosynthetic process”, and “Oxoacid metabolic process” (Fig. 4h). JA production is a carboxylic acid metabolic process and interactors linked to these GO terms include the JA synthesis enzymes CYP74A and LOX2 (Bell et al. 1995; Laudert et al. 1996). The top five most enriched cellular component GO terms for the THE1_Δkinase_ interactors are “Chaperonin-containing T-complex”, “Peroxisome”, “Microbody”, “Chloroplast stroma”, and “Plastid stroma” (Fig. S8). The results indicate that kinase domain removal shifts the interactome in comparison to wild-type THE1 towards proteins apparently involved in JA-related metabolic processes.

For THE1_K636E_, we found 43 significantly enriched interactors (Fig. 4i; Data S1). The most significant interactors for THE1_K636E_ include AT1G65720, RECEPTOR-LIKE KINASE 7 (RLK7) (Pitorre et al. 2010), LOX2 (Bell et al. 1995), PHLOEM PROTEIN 2-LIKE A1 (PP2A1) (Beneteau et al. 2010), and GDPDL3 (Fig. 4c). The most enriched biological process GO terms for THE1_K636E_ interactors are “Response to external (biotic) stimulus”, “Response to other organism”, and “Biological response involved in interspecies interaction between organisms”. The five most significant cellular component GO terms for THE1_K636E_ are “Cell-cell junction”, “Plasmodesma”, “Cell junction”, “Symplast”, and “Anchoring junction” (Fig. S8). The GO terms and subcellular localization are the same for THE1_K636E_ and wild-type THE1 (Fig. 3d and e). This probably indicates that the K636E mutation reduced the number of interactors we observed but does not alter the dominant biological processes and subcellular components associated with the interactors.

Taken together, we identified distinct sets of specific interactors for the THE1 variants THE1_G37D_, THE1_ΔECD_, THE1_Δkinase_, and THE1_K636E_. Our results suggest misfolding of THE1_G37D_ and its retention in the ER. The K636E mutation of the THE1 kinase domain alters the interactors in comparison to the wild-type THE1, and the complete absence of the kinase domain shifts the interactors towards an association with JA metabolism.

### Stress treatments identify stress-specific THE1 interactors

THE1 is required for the response to ISX-induced cell wall damage and associated with the osmotic stress response (Hématy et al. 2007; Bacete et al. 2022). We hypothesized that particular interactions could be stress-specific and performed IP-LC-MS/MS after exposing seedlings to half-strength Murashige and Skoog (½ MS) medium with DMSO (solvent control), the cellulose biosynthesis inhibitor ISX, the osmoticum sorbitol, or a combined ISX-sorbitol treatment (ISX-induced responses can be suppressed by osmoticum (Hamann et al. 2009; Wormit et al. 2012)) (Fig. 5a). The four-hour period was chosen because THE1-modulated hyperosmotic stress-induced ABA production peaks at this timepoint (Bacete et al. 2022). Furthermore, ISX-induced responses, such as the pattern-triggered immunity-like transcriptional response, start to be activated at four hours of treatment (Zhai et al. 2024). Moreover, we characterized six-day-old seedling roots expressing THE1-GFP after four hours of DMSO, ISX, sorbitol, and ISX-sorbitol treatment (Fig. S9). While no obvious changes in the localization of THE1-GFP were observed, cells in the root tip began to swell after four hours of ISX or combined ISX-sorbitol treatment, confirming that ISX leads to CWI impairment at four hours.

**Figure 5.**
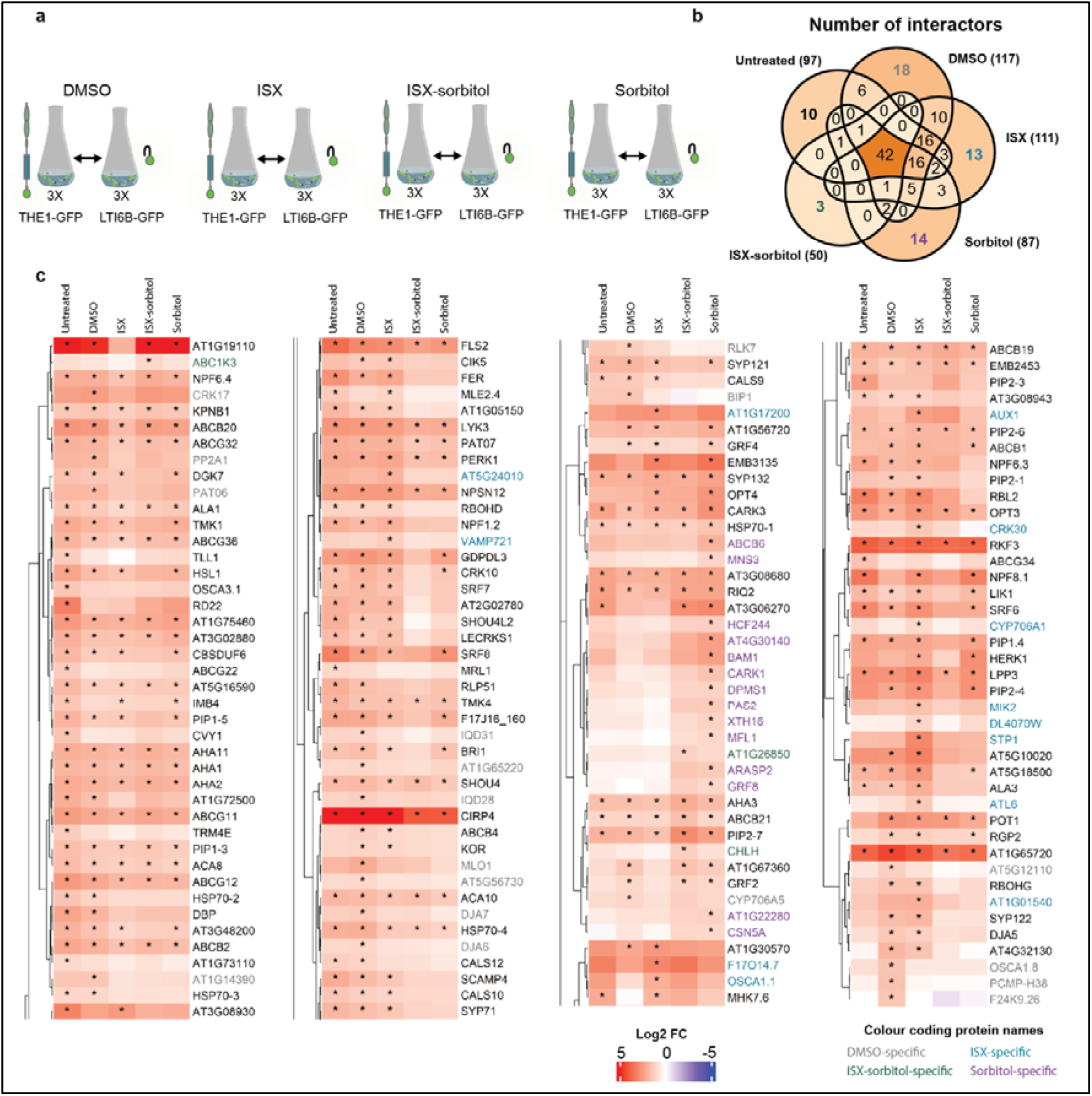
Putative condition-specific THE1 interactors as identified by immunoprecipitation-liquid chromatography tandem mass spectrometry (IP-LC-MS/MS). **a)** IP-LC-MS/MS was performed on liquid-grown seven-day-old seedlings expressing THE1-GFP and LTI6B-GFP (n=3). Seedlings were treated for four hours with medium containing DMSO, ISX (600 nM), combined ISX-sorbitol treatment (600 nM and 300 mM, respectively), or sorbitol (300 mM). **b)** Venn diagram comparing the number and overlap of significant THE1 interactors between the different treatments. The total interactor number per treatment type is indicated in parentheses. Interactors identified in untreated samples correspond to Fig. 3b. **c)** Heatmap showing the log_2_ fold-change (FC) in which FC is calculated as FC = (protein group intensity THE1-GFP/protein group intensity LTI6B-GFP). Rows were clustered based on Pearson’s correlation coefficient. Asterisks indicate significantly enriched proteins, as identified using the R-package *limma* (Ritchie et al. 2015) with Strimmer’s Q-value false discovery rate (FDR) correction. Proteins with a fold-change (FC) >2 and adjusted P-value <0.05 were considered significantly enriched. The heatmap was split into four parts to match figure size limits. The names of interactors which were only significantly enriched in one specific treatment condition were color-coded in grey (DMSO), blue (ISX), green (ISX-sorbitol), and purple (sorbitol).

We found 117 putative interactors after treatment with DMSO, 111 with ISX, 87 with sorbitol, and 50 with combined ISX-sorbitol (Fig. S10; Data S1). In the same experiment, 97 interactors were detected for untreated samples for which the medium was not replaced (corresponds to Fig. 3b). Next, we compared the interactome of THE1 in stress-treated samples to the interactome of THE1 in untreated samples (Fig. 5b). This identified a set of 42 interactors that were enriched in all samples, including ACA8 and -10, AHA1, -2, -3, and -11, ABCB2, -19, and 21, ABCG32 and -36, FLS2 and CIRP4. These interactors may perform stress-independent functions or are components of pre-assembled signaling platforms activated post-translationally upon stress.

To identify the treatment-specific interactors, we visualized the results in a heatmap (Fig. 5c). We considered the differences in interactors between untreated and DMSO-treated samples. DMSO was used as a solvent for ISX and was added to all treatments to allow for treatment-specific comparisons. The difference between untreated and DMSO-treated samples is not only the addition of DMSO, but also replenishment of nutrients and sucrose by exchanging the growing medium. Ten interactors were specifically enriched in untreated conditions, including PIP2-3 (Johanson et al. 2001), ABCG22 (Johanson et al. 2001), ABCG34 (Khare et al. 2017), and OSCA3.1 (Zhang et al. 2018b). A group of 36 interactors was enriched in DMSO-treated samples but not in untreated samples. This difference in interactors suggests an effect on the *A. thaliana* seedlings by the presence of DMSO alone or nutrient/sugar replenishment resulting from refreshing the growing medium. Examples are KORRIGAN (KOR) (Sato et al. 2001), BIP1 (Koizumi 1996), RESPIRATORY BURST OXYDASE HOMOLOG G (RBOHG) (Torres and Dangl 2005), PROTEIN S-ACYLTRANSFERASE 6 (PAT06) (Batistič 2012), RLK7 (Pitorre et al. 2010), and MILDEW RESISTANCE LOCUS O 1 (MLO1) (Devoto et al. 2003). These interactors may play a role in the response to DMSO, sucrose, or shifting nutrient levels.

Interestingly, several proteins were specifically enriched in response to cell wall damage or hyperosmotic stress: 13 for ISX, 14 for sorbitol, and three for ISX-sorbitol. ISX-specific interactors include the leucine-rich repeat receptor kinase MALE DISCOVERER 1-INTERACTING RECEPTOR-LIKE KINASE 2 (MIK2) (Wang et al. 2016; Van der Does et al. 2017), the RING-type ubiquitin ligase ARABIDOPSIS TOXICOS EN LEVADURA 6 (ATL6) (Salinas-Mondragón et al. 1999; Sato et al. 2009), and VESICLE-ASSOCIATED MEMBRANE PROTEIN 721 (VAMP721) (Uemura et al. 2004; Kwon et al. 2008). Amongst the sorbitol-specific interactors are CYTOSOLIC ABA RECEPTOR KINASE 1 (CARK1) (Zhang et al. 2018a) and GENERAL REGULATORY FACTOR 8 (GRF8) (Camoni et al. 2018). Interactors exclusive to ISX-sorbitol co-treated samples are MAGNESIUM-CHELATASE SUBUNIT H (CHLH) (Mochizuki et al. 2001), ACTIVITY OF BC1 COMPLEX KINASE 3 (ABC1K3) (Lundquist et al. 2013), and the methyltransferase homolog AT1G26850 (Mouille et al. 2007). We note that the interaction of THE1 with the latter proteins should be critically assessed; CHLH and ABC1K3 have a predicted plastid localization (Mochizuki et al. 2001; Lundquist et al. 2013) (Data S1), while THE1 is located at the plasma membrane (Fig. 1). Three interactors overlap between ISX- and sorbitol-treated *A. thaliana*: HERK1 (Guo et al. 2009), OLIGOPEPTIDE TRANSPORTER 4 (OPT4) (Osawa et al. 2006), and EMBRYO DEFECTIVE 3135 (EMB3135) (Meinke 2020). This overlap suggests a role of these proteins in different CWI-impairing processes.

Overall, our interaction study identified multiple novel candidates that interact with THE1 specifically during exposure to cell wall damage or hyperosmotic stress. Simultaneously, many THE1 interactors are found both in ISX- and sorbitol-treated samples, suggesting that *A. thaliana* uses correlated or intersecting signaling pathways to maintain CWI.

## Discussion

THE1 is an evolutionary conserved RK required for CWI maintenance (Hématy et al. 2007; Zwartkruis et al. 2026). It activates responses to impaired cellulose biosynthesis, acts as a positive regulator of JA production, acts as a negative regulator of ABA production in response to hyperosmotic stress, and modulates cell wall stiffness (Hématy et al. 2007; Engelsdorf et al. 2018; Bacete et al. 2022). The *the1-1* and *the1-4* alleles have opposing effects on the strength of responses to cellulose biosynthesis inhibition (Merz et al. 2017; Bacete et al. 2022), making them perfect tools to study the mode of action of THE1 in CWI maintenance, where knowledge currently remains limited (Zwartkruis et al. 2026). To initiate a functional analysis of THE1, we generated a set of transgenic THE1-GFP lines in the *the1-6* (S53STOP; Merz et al. 2017) *A. thaliana* background (Fig. 1).

One of the reasons why biochemical characterization of THE1 remained understudied may be due to the molecular cloning of *THE1*. Our initial attempts to clone a full-length version of *THE1* or *THE1_G37D_*resulted in constructs with undesirable mutations or truncated sequences (in various *Escherichia coli* strains), or no *E. coli* colonies at all. However, clones of kinase-mutated and kinase-dead THE1 (like THE1_K636E_ and THE1_D634N_, Fig. S2) could be obtained, suggesting cloning success may depend on kinase activity. Interestingly, *THE1*_Δ*ECD*_ could be cloned, despite the kinase-domain being present. To obtain a full-length *THE1 and THE1_G37D_*, we cloned the ORFs in two parts in entry plasmids, confirmed their integrity by sequencing and assembled them into one destination plasmid using a GreenGate reaction (Lampropoulos et al. 2013). These plasmids were directly transformed into *Agrobacterium tumefaciens* used to generate stable transgenic *A. thaliana* lines, and sequencing confirmed integrity of full-length sequences *in planta*. Overall, our alternative cloning strategy allowed for the generation of unmutated THE1-GFP and THE1_G37D_-GFP *A. thaliana* lines. The alternative cloning strategy used here may be applied to clone other *CrRLK1Ls* and other genes that face similar cloning challenges in the future. Together with GFP-fusion lines of THE1_K636E_, THE1_Δkinase_ and THE1_ΔECD_-GFP, we generated a set of THE1-GFP variants allowing functional characterization of THE1 by localization and interaction studies.

Using *THE1_pro_:THE1-GFP A. thaliana*, we generated an *in vivo* map of THE1 localization. THE1-GFP localized ubiquitously in roots and shoots of six-day old seedlings, with the highest abundance in the root tip and root vasculature and lowest abundance in the basal root region (Fig. 1b; Fig. S5). A previous study used a *THE1_pro_:*β*-GLUCURONIDASE* construct and detected *THE1_pro_* activity in the same regions where we observe the protein except for the distal meristematic zone, where reporter activity was absent (Hématy et al. 2007). Another reporter line of *THE1_pro_*activity (*THE1_pro_:YFP*) showed expression in the root tip of six-day-old seedlings, in line with our observations of protein localization (Bacete et al. 2022). The ubiquitous expression of THE1 suggests that it performs a function in many different cell types. At the subcellular level, we showed that THE1-GFP localizes to the plasma membrane (Fig.1c and 1d), in line with earlier reports of plasma membrane localization of THE1-GFP expressed under control of *35S_pro_* in hypocotyl cells (Hématy et al. 2007). The localization of RKs can change upon (stress-induced) activation or ligand binding (Di Rubbo and Russinova 2012). For example, FER is internalized after treatment with its ligand RALF1 (Rößling et al. 2024). We did not observe changes in localization of THE1-GFP after four hours of ISX, sorbitol or combined ISX-sorbitol treatments (Fig. S9). Future studies may test how THE1 localization changes upon treatment with RALFs such as RALF34, or perform short-term stress treatments to identify activators of THE1 and understand temporal dynamics better. THE1_K636E_-GFP and THE1_Δkinase_ localize to the plasma membrane (Fig. 1c and d), indicating that the kinase domain is not required for plasma membrane localization. In contrast, THE1_G37D_-GFP and THE1_ΔECD_-GFP were found to localize mainly intracellularly, and to a limited extent or not at all to the plasma membrane (Fig. 1c and d; Fig. S6), indicating that mutations in the THE1 ectodomain may alter THE1 plasma membrane localization.

THE1_G37D_ is encoded by the *the1-1* loss-of-function allele (Hématy et al. 2007). The glycine residue mutated in this allele is conserved in the *Cr*RLK1L family (Zwartkruis et al. 2026) but the specific cause for the loss of function was unknown (Gonneau et al. 2018). While THE1-GFP localizes to the cell periphery, THE1_G37D_-GFP displays intracellular localization (Fig. 1c and d; Fig. S6). Moreover, in contrast to full length THE1, THE1_G37D_ interacts with molecular components of the ER-localized protein quality control mechanism (Kleizen and Braakman 2004) such as BIP1 and -2 (Koizumi 1996; Martínez and Chrispeels 2003), UGGT (Blanco-Herrera et al. 2015), ERDJ3B (Shen and Hendershot 2005), and CNX1 (Liu et al. 2017) (Fig. 4g and h). This evidence suggests that the *the1-1* mutant is a loss-of-function mutant because THE1_G37D_, encoded in *the1-1,* does not fold correctly and is therefore retained in the ER. The BRI1 loss-of-function mutant encoded in the *bri1-5* allele (C69Y mutation) is another example of an RK with an ectodomain point mutation that showed ER retention and interaction with CNX1 and BIPs (Hong et al. 2008). An example of an animal RK with similar defects is provided by F409V mutation of the insulin receptor (F382V in old residue numbering nomenclature used in the reference), which also causes ER retention and interaction with BIP (Accili et al. 1989, 1992). In FER, the corresponding G41S mutation causes failure of root hair formation at elevated temperatures (Kim et al. 2021). In contrast to the THE1 G37D mutation, FER G41S causes normal localization at room temperature but reduced the presence of FER in the apical root tip at elevated temperatures (Kim et al. 2021). A possible explanation for this temperature-dependent effect is that in FER two additional glycine residues downstream of G41 (absent in THE1) may enable correct folding at lower temperatures (Zwartkruis et al. 2026). The *the1*-*2* allele (E150K mutation) is a second *the1* loss-of-function allele with a single ectodomain-localized amino acid substitution (Hématy et al. 2007). We propose that THE1_E150K_ could be tested for ER localization in the future to establish if its loss of function stems from the same cause or may be due to other causes such as ligand binding defects. Overall, our results suggest that the *the1-1* allele is a loss-of-function mutant because of protein folding defects, answering a 20-year-old research question (Hématy et al. 2007).

The malectin-like domain-lacking THE1_ΔECD_ also showed mainly intracellular localization (Fig. 1c and d). A similar variant of FER lacking the ectodomains localized to the peripheral ER (Qin et al. 2026), and THE1_ΔECD_ may also localize there. A co-localization study using ER marker proteins such as the V-ATPASE ASSMBLY FACTOR AtVMA21a (Neubert et al. 2008) may be performed to test putative ER localization in the future. Yet, we observed that not a single quantified THE1_ΔECD_ interactor that has predicted ER localization (Data S1). Further characterization of THE1_ΔECD_-GFP localization and the consequences of absence of malectin-like domain for THE1-mediated responses is needed before any functional relationships between THE1_ΔECD_ and its interactors can be confidently suggested.

The full-length THE1 localizes to the plasma membrane (Fig. 1c and d; Fig. S6), and we identified putative interactors of THE1 in control conditions (Fig. 3; Table 1), and in conditions in which CWI is impaired (Fig. 5). The interactors seem to reflect the diverse processes in which THE1 plays a role (Zwartkruis et al. 2026), including the response to cellulose biosynthesis inhibition for which THE1 was originally characterized (Hématy et al. 2007). Two regulators of cellulose synthesis may be especially relevant: GDPDL3 (also known as SHAVEN3 (Hayashi et al. 2008)) and SHOU4 (Polko et al. 2018). GDPDL3 is a glycosylphosphatidylinositol-anchored protein localized in the outer plasma membrane leaflet initially studied for its role in root hair growth (Parker et al. 2000; Jones et al. 2006). Double mutants of *gdpdl3* and its paralog *gdpdl4* display reduced cellulose content, ruptured root hairs, expansion of the hypocotyl epidermis, and ectopic lignin deposition (Hayashi et al. 2008; Yeats and Somerville 2016). SHOU4 is a plasma membrane-localized protein which acts as a negative regulator of CELLULOSE SYNTHASE COMPLEX exocytosis (Polko et al. 2018). When the function of both SHOU4 and its homolog SHOU4-LIKE are impaired, plants have increased levels of amorphous cellulose and root hairs fail to emerge (Polko et al. 2018). We are not aware of root hair phenotypes in *THE1* mutants, but root hair defects are commonly observed in loss-of-function *fer* and *eru* mutants (Duan et al. 2010; Bai et al. 2014; Schoenaers et al. 2018). SHOU4 regulates the number of plasma membrane-cell wall attachment sites and is important for both biotic and abiotic stress responses (Wang et al. 2023; Rui et al. 2026). GDPDL3 and SHOU4 are exciting future study subjects to understand the function of THE1 in response to cellulose biosynthesis inhibition and in stress responses.

THE1 and several of the potential interactors have been associated with the ABA-mediated stress responses (Bacete et al. 2022). We found KPNB1 and its homolog AT3G08943 (59.9% amino acid sequence identity) as putative interactors. KPNB1 is an importin β protein, which negatively regulates drought tolerance and ABA signaling (Luo et al. 2013). Importins are nuclear transport receptors that mediate shuttling of proteins and other cargo into the nucleus (Xiong et al. 2021). These proteins may link THE1 to transcriptional regulation of ABA mediated processes. The role of THE1 in ABA responses can also depend on its interaction with DGK7 (Table 1), which produces the lipid second messenger PA (Gómez-Merino et al. 2004, 2005). One of the many physiological roles of PA is that it can inhibit ABSCISIC INSENSITIVE 1 (Koornneef et al. 1984; Zhang et al. 2004; Testerink and Munnik 2011), which is a phosphatase that negatively regulates ABA responses (Leung et al. 1994; Meyer et al. 1994). Other ABA-related interactors are CARK1, which interacted with THE1 in sorbitol treatment, and its homolog CARK3, that interacted independent of treatment type (Fig. 5). CARK1 and CARK3 are RECEPTOR-LIKE CYTOPLASMIC KINASE proteins, which are often directly downstream of RKs in signaling cascades (Boisson-Dernier et al. 2015; Bender and Zipfel 2023). CARK1 phosphorylates ABA receptor proteins, providing a link to ABA signaling (Zhang et al. 2018a). CARK1 and CARK3 were recently shown to phosphorylate AHA2 and may also mediate THE1-dependent regulation of extracellular pH (Li et al. 2026b).

The regulation of extracellular pH appears to be a conserved role of members in the *Cr*RLK1L family and their RALF ligands (Pearce et al. 2001; Haruta et al. 2014; Gonneau et al. 2018; Schoenaers and Vissenberg 2025). We found AHA1, -2, -3, and -11 as putative interactors of THE1 (Fig. 3a and b). Extracellular pH determines the activity of cell wall loosening proteins known as expansins to allow cell expansion (Cosgrove 2024a, 2024b). THE1 has been connected to modulation of cell expansion and cell wall stiffness (Hématy et al. 2007; Bacete et al. 2022; Alonso Baez et al. 2026). We propose that THE1 regulates the activity of AHA1, -2, -3, and -11, leading to adaptive modifications of pH and cell wall structure, and future studies could elucidate how THE1 regulates these AHAs specifically. Interactions between *Cr*RLK1L family members were hypothesized to be relevant in determining *Cr*RLK1L-RALF signaling (Gonneau et al. 2018; Schoenaers and Vissenberg 2025). Recently, the THE1 ectodomain was observed to co-precipitates with the ectodomains of FER, ANJ, and HERK1 in a co-IP experiment (Li et al. 2026a). We repeatedly find both FER and CURVY1 as THE1 interactors (Fig. 2a and b; Table 1). Reciprocally, THE1 was also identified as a FER interactor in IP-LC-MS/MS studies (Leicher et al. 2026). Interestingly, HERKULES1, another *Cr*RLK1L involved in petiole elongation and salt stress responses (Guo et al. 2009; Gigli-Bisceglia et al. 2022), was detected as a putative THE1 interactor only in ISX-and sorbitol-treated samples (Fig. 5c). While the functional relevance of these interactions remains unclear, the selectivity suggests they are regulated in a context-dependent manner.

Furthermore, the AHAs may also be among those interactors by which THE1 activates cell wall damage-induced immune responses (Denness et al. 2011; Engelsdorf et al. 2018). It has long been recognized that AHA activity determines the strength of plant immune and wounding responses (Schaller and Oecking 1999; Zhai et al. 2026). A specific example is that JA production and the duration and amplitude of the slow wave potential that follow wounding depends on AHA1 (Kumari et al. 2019). The interactor ABCG34 (Fig. 5c) may also be involved in the immune responses mediated by THE1 (Engelsdorf et al. 2018), because secretion of the antimicrobial compound camalexin depends on *ABCG34* expression levels (Ahuja et al. 2012; Khare et al. 2017). Interestingly, ISX-induced camalexin production is also THE1-dependent (Morton et al. 2026). The data we present suggests that a physical interaction between ABCG34 and THE1 may be relevant in determining camalexin production. Further interactors cooperating with THE1 to elicit ISX-induced immune responses may also include MIK2 (Wang et al. 2016; Van der Does et al. 2017), ATL6 (Salinas-Mondragón et al. 1999; Sato et al. 2009), and VAMP721 (Uemura et al. 2004; Kwon et al. 2008), which we observed specifically as THE1 interactors after ISX treatment (Fig. 5c). MIK2 is a RK that has diverse functions including mediating the ISX-induced pattern triggered immunity response (Van der Does et al. 2017; Zhai et al. 2024). MIK2 probably acts downstream of THE1, because ISX-induced expression of the MIK2 ligand SERINE RICH ENDOGENOUS PEPTIDE 18 depends on THE1 (Zhai et al. 2024). ATL6 is a ubiquitin ligase that is transcriptionally induced by cellulase treatment (Salinas-Mondragón et al. 1999; Sato et al. 2009). FER can phosphorylate ATL6 to regulate responses to carbon/nitrogen nutrient ratios (Xu et al. 2019), providing an example that *Cr*RLK1Ls can phosphorylate ubiquitin ligases. Our results indicate that THE1 may be involved in regulation of ATL6 as well. VAMP721 and its homolog VAMP722 are vesicle-associated membrane proteins and function in focal secretion at sites of fungal attack (Kwon et al. 2008). Fungi secrete cell wall degrading enzymes or can penetrate the cell wall by force (Kubicek et al. 2014; Bronkhorst et al. 2021). THE1 may sense locally impaired CWI and recruit VAMP721 as a response to invading pathogens.

Information about the specific role of THE1 as a positive regulator of JA production can be deduced from results of the *the1*-*4* mutant, encoding a version of THE1 lacking the kinase domain (Guo et al. 2009; Merz et al. 2017). We observed increased JA levels in *the1-4* seedlings relative to Col-0 plants in both ISX- and DMSO-treated seedlings (Fig. 2a and b), similarly to previous reports (Engelsdorf et al. 2018; Bacete et al. 2022). However, it is not known how a kinase-lacking version of THE1 causes increased JA production (Merz et al. 2017). One explanation for the increased JA levels observed in the *the1*-*4* mutant would be based on an interaction with FER. *A. thaliana* seedlings with the *fer*-*5* knock-down allele show increased JA production already in control conditions (Duan et al. 2010; Engelsdorf et al. 2018). Co-IP experiments recently showed that the THE1 ectomain co-precipitates with the FER ectodomain (Li et al. 2026a). An interaction between FER and a kinase-lacking variant of THE1 may lead to (partial) loss of function of FER, lifting its inhibitory action on JA production. Interestingly, we also observed slightly elevated levels of JA in seedlings expressing the kinase-impaired THE1_K636E_-GFP construct (Fig. 2a and b). Hence, it seems that lack of a THE1 kinase domain or impaired THE1 kinase activity causes increased JA levels, and we hypothesize that this may be caused by changes in the interactions between THE1 and FER. The potential interactors we detected and quantified for THE1_Δkinase_ (corresponding to *the1-4*) are mostly enzymes induced by JA or involved in JA biosynthesis, such as LOX2 (Bell et al. 1995), TAT3 (Riewe et al. 2012; Liu and Avramova 2016), CYP96A4 (Huang et al. 2024), CYP74A (Laudert et al. 1996), and MDAR3 (Sasaki-Sekimoto et al. 2005) (Fig. 4c and d). Some other THE1_Δkinase_ interactors are involved in organelle fission. The interactor ARC5 is involved in chloroplast fission (Gao et al. 2003), while DRP3A and its homolog DRP3B are involved in peroxisome and mitochondrion fission (Fujimoto et al. 2009). These interactors are not well-established interactors for receptor-like proteins (the architecture of THE1_Δkinase_ is similar to a receptor-like protein (Jeong et al. 1999)) or RKs (Bender and Zipfel 2023; Huang and Joosten 2025). Moreover, the interactors are not enriched for the “Plasma membrane” GO term while we observe THE1_Δkinase_ to localize to the plasma membrane. This raises the question to what extent the observed THE1_Δkinase_ interactors are *bona fide* interactors, or rather enriched proteins in this mutant background which already showed elevated JA levels in DMSO-treated controls (Fig. 2a).

The role of the THE1 kinase is further demonstrated by results of THE1_K636E_. This variant, introduced as THE1_K636E_-GFP in *the1-6* background, restored ISX-induced JA production (Fig. 2b) and displayed higher-than-wildtype levels of JA production in control conditions (Fig. 2a). K636 is the catalytic loop lysine in the HRD-X-K motif and is strongly conserved in serine/threonine kinases (Hanks et al. 1988; Kornev et al. 2006), where it stabilizes the transition state of phosphoryl transfer (Cheng et al. 2005). Our data suggests that K636E mutation of THE1 has a limited effect on autophosphorylation activity but abolishes transphosphorylation activity (Figs. S1 and S2). Mutation of the HRD-X-K motif lysine in the yeast cAMP-dependent protein kinase homolog TAKASHI’S PROTEIN KINASE 1 resulted in a 99.7% decrease in substrate phosphorylation (Gibbs and Zoller 1991), underscoring the importance of this lysine residue in serine/threonine kinases for kinase activity. THE1D634N, harboring a mutation in the HRD motif commonly used to abolish kinase activity (Hanks et al. 1988; Kimura et al. 2020), showed no autophosphorylation activity in the kinase assay (Fig. S2). We recommend THE1_D634N_ as a kinase-dead THE1 variant in addition to the kinase-mutated THE1_K636E_ in the future. A kinase-dead FER rescued the reduced ovule fertilization phenotype of the *fer-1* loss-of-function mutant (Huck et al. 2003; Escobar-Restrepo et al. 2007; Kessler et al. 2015). Our results provide another example that kinase activity of *Cr*RLK1Ls is not always required for their function.

Interactors present in THE1 but not in THE1_K636E_ may be interactors that depend on kinase activity. CIRP4 is a strongly enriched THE1 interactor (Fig. 3a and b) that is absent in THE1_K636E_ (Fig. 4i). Based on its homology with the characterized ubiquitin ligase CIRP1 (54% amino acid identity) (Yang et al. 2025), CIRP4 seems to be a ubiquitin ligase. The absence of CIRP4 among the THE1_K636E_ interactors may indicate that CIRP4 specifically interacts with a functional THE1 kinase. We hypothesize that THE1 phosphorylates CIRP4, similarly as FER phosphorylates ATL6 (Xu et al. 2019). The CIRP4 homolog CIRP1 regulates levels of reactive oxygen species by direct ubiquitination of catalases (Yang et al. 2025), which are enzymes that break down the reactive oxygen species H_2_O_2_ (Loew 1900; Frugoli et al. 1996). We propose that CIRP4 may perform a similar role to CIRP1, and that THE1 could regulate levels of reactive oxygen species by this mechanism. Regulation of H_2_O_2_ levels can play a signaling role, as osmotic stress was recently shown to result in H_2_O_2_-rich nanodomains where the molecular switch RHO OF PLANTS 6 accumulates (Poitout et al. 2026). H_2_O_2_ also has a role in determining the mechanical properties of cell walls by chemically crosslinking phenolic compounds (including lignin) (Schopfer 1996; Xiong et al. 2015), providing a possible mechanism by which THE1 may influence cell wall stiffness. Alternatively, CIRP4 may be an ubiquitin ligase that ubiquitinates THE1 itself, since THE1 is described to be post-translationally modified by ubiquitination (Grubb et al. 2021; Zwartkruis et al. 2026). Another recurrent interactor, AT1G19110, is related to CIRP4 (based on phylogeny) and may also have ubiquitin ligase activity (Wang et al. 2024). Future studies should address the possible interaction between CIRP4 and THE1 in the context of either reactive oxygen species, proteostasis, or ubiquitin signaling.

A limitation of IP-LC-MS/MS-based interaction studies is that transient and indirect interactors may not co-precipitate and are therefore not detected. This could explain why GEF4, which was previously shown to interact with THE1 (Qu et al. 2017), is not detected as significantly enriched in the THE1 interactome we identified. Future studies can use cross-linking or proximity labeling to identify also more transient THE1 interactors (Özmen et al. 2025; Bai et al. 2026). The C-terminal fusion of GFP to THE1 could sterically hinder interactors from binding and exclude them from detection in IP-LC-MS/MS, which may be addressed by repositioning the GFP tag or employing a smaller tag such as the HA tag or FLAG tag (Kimple et al. 2013). We mainly identified transmembrane- or intracellularly-localized proteins as THE1 interactors (Fig. 3f), possibly because extracellular proteins remain bound to the cell wall fraction and do not co-precipitate with the THE1-GFP bait (Albenne et al. 2013). Proteins embedded in the cell wall matrix could be isolated using specialized methods (Feiz et al. 2006; Albenne et al. 2013), and subsequently combined with the soluble fraction to increase the chance of detecting extracellular interactors. Moreover, extracellular proteins are often glycosylated, potentially hindering trypsin digestion and detection during MS (Albenne et al. 2013). Enzymatic deglycosylation of the samples before MS may remove this limitation (San Clemente and Jamet 2022). THE1 has a predicted N-glycosylated ectodomain (Zwartkruis et al. 2026), and we observed that few peptides in the IP-LC-MS/MS results map to the ectodomain, probably as a result of resistance to trypsin digestion. THE1 may not show up as strongly enriched in the IP-LC-MS/MS data of THE1_Δkinase_ because of trypsin indigestibility (Fig. 4g). Another example of proteins possibly not detected due to their glycosylation are members of the LORELEI (LRE) protein family, which are the canonical co-receptors of *Cr*RLK1Ls (Li et al. 2015; Xiao et al. 2019; Noble et al. 2022; Zwartkruis et al. 2026). A more targeted approach, such as co-IP experiments, may be performed to identify if the LRE family members interact with THE1. Finally, enrichment of proteins in an IP-LC-MS/MS experiment is not a proof of direct interaction. Orthogonal validation by methods such as yeast two-hybrid, bimolecular fluorescence complementation or Förster resonance energy transfer is required in the future to confirm direct interactions of individual candidates with THE1 discovered here (Fields and Song 1989; Strotmann and Stahl 2022).

This work presents the results from a systematic analysis of the THE1 interactome. We identified candidates by which THE1 may regulate cell wall stiffness, JA production, and hyperosmotic stress-induced ABA levels, and explained the mechanism by which G37D mutation of THE1 causes loss of function. Our study lays the foundation for future works to identify the hierarchy and functional relationship between THE1, its interactors, and their role in CWI maintenance.

## Supporting information

Interactors

Protein intensities

## Acknowledgements

The authors thank all members of the Hamann lab for feedback on the project and the manuscript. The authors thank Martijn Vandegehuchte and Debora Gasperini for fruitful discussions about the project. We are grateful to Mari Fjøsne for help in the lab.

## Author contributions

S.T.W.Z., W.H., T.T., and T.H. designed the research. S.T.W.Z., W.H., T.T., N.B., Z.B., A.Z., M.W, L.H., V.J.K., D.A., L.A.B., and L.H. performed experiments. G.M. provided the kinase structural model. S.T.W.Z, Z.B., and A.S. performed data analysis. T.H. obtained funding. S.T.W.Z wrote the original draft with input from W.H and T.H.; all authors provided feedback and approved the final manuscript.

## Supplementary material

Supplementary material is available online.

## Data availability

The mass spectrometry proteomics data have been deposited to the ProteomeXchange Consortium via the PRIDE101 partner repository with the dataset identifier XXX.

