## Supplemental information for "Mapping the THESEUS1 interactome links cell wall integrity and stress signalling in Arabidopsis"

**Table S1:** Primers used in this study

**Figure S1**: Structural model of a THESEUS1 kinase-domain dimer with the activation loop of one protomer bound in the active site of the other.

**Figure S2**: Kinase activity assay showing the kinase activity of THE1_K636E_

**Figure S3**: Representative images of rosettes of four-week-old *Arabidopsis thaliana* wildtype plants, *THE1* mutants, and *THE1-GFP* transgenic lines

**Figure S4**: Representative images of five true leaves starting from the sixth true leaf in rosettes of four-week-old *A. thaliana* wildtype plants, *THE1* mutants, and *THE1-GFP* transgenic lines.

**Figure S5**: Overview of *THE1_pro_:THE1-GFP* localization in a six-day-old *A. thaliana* seedling.

**Figure S6**: Plasmolyzed epidermal cells in the elongation zone of six-day-old *A. thaliana* seedlings expressing THE1-GFP variants.

**Figure S7**: InterProScan annotation of the *A. thaliana* protein of unknown function AT1G65720.

**Figure S8**: Significantly enriched cellular component gene ontology (GO)-terms associated with THE1 variant interactors.

**Figure S9**: Localization of THE1-GFP in six-day-old *A. thaliana* seedling roots after four-hour treatment with DMSO, ISX, Sorbitol, or combined ISX-sorbitol.

**Figure S10:** Putative THE1 interactors identified by immunoprecipitation–liquid chromatography–tandem mass spectrometry after four-hour treatment with DMSO, ISX, sorbitol, or combined ISX-sorbitol.

**Figure S11**: Sequencing of constructs containing *THE1*, *THE1_G37D_*, *THE1_K636E_*

**Figure S12**: Sequencing of construct containing *THE1_Δkinase_*

**Figure S13**: Sequencing of construct containing *THE1_ΔECD_*

**Figure S14**: Sequencing of *THE1* and *THE1_G37D_* from *A. thaliana* stable lines transformed with *THE1-GFP* or *THE1_G37D_-GFP*.

**Table S1** Primers used in this study. Underlined: restriction enzyme recognition site, bold and italics: overhang for GreenGate system (Lampropoulos et al. 2013). atta/taat additional nucleotides allow for better restriction digest of PCR product.

| **Primer Name** | **Primer Sequence** | **Primer Purpose** |
| --- | --- | --- |
| THE1pro_BsaI_GGA_FW | attaGGTCTCt***ACCT***aaaacttacatataattaacttctcaaaatgttttatc | cloning THE_pro_ in pGGA000, "A" overhang for GreenGate system |
| THE1pro_BsaI_GGB_REV | taatGGTCTCt***TGTT***tttggatttttagtaagaaccctaaaacc | cloning THE_pro_ in pGGA000, "B" overhang for GreenGate system |
| 8A-linker-GFP_BsaI_GGD_FW | attaGGTCTCt***TCAG****GG*GCAGCTGCAGCTgcggccgctgcaatgg | cloning eGFP into pGGD000 with 8-alanine-linker, "D" overhang for GreenGate system |
| 8A-linker-GFP_BsaI_GGE_REV | taatGGTCTCt***GCAG***ctacttgtacagctcgtccatg | cloning eGFP into pGGD000 with 8-alanine-linker, "E" overhang for GreenGate system |
| THE1_pIBV5_TOPO_FW | AAAATGGTGTTCACAAAATCATTACTTGTTC | cloning THE1_Δkinase_ in pIB/V5-His TOPO |
| SAIL_LB1 | GCCTTTTCAGAAATGGATAAATAGCCTTGCTTCC | cloning THE1_Δkinase_ in pIB/V5-His TOPO |
| THE1_noBsaI_FW | ACCATTTAGTggactcTCTCTGCTTG | site-directed mutagenesis to remove the *Bsa*I restriction enzyme recognition site in *THE1* |
| THE1_noBsaI _REV | GTTGAAGGGTTTAACGCC | site-directed mutagenesis to remove the *Bsa*I restriction enzyme recognition site in *THE1* |
| THE1_BsaI_GGC_FW | attaGGTCTCt***GGCT***ATGGTGTTCACAAAATCATTAC | cloning THE1_Δkinase_ in pGGC000 (incl. signal peptide), "C" overhang for GreenGate system |
| the1-4_BsaI_GGD_REV | taatGGTCTCt***CTGA***ATTTGTCTGAGAGAGTCC | cloning THE1_Δkinase_ in pGGC000, "D" overhang for GreenGate system |
| THE1.pt1_BsaI_GGB_FW | attaGGTCTCt***AACA***ATGGTGTTCACAAAATCATTAC | cloning THE1- and THE_G37D_-pt1 in pGGB000 (incl. signal peptide), "B" overhang for GreenGate system |
| THE1.pt1_BsaI_GGC_REV | taatGGTCTCt***AGCC***ATACCTTGTTCTGAT | cloning THE1- and THE_G37D_-pt1 in pGGB000 (incl. signal peptide), "C" overhang for GreenGate system |
| THE1.pt2_BsaI_GGC_FW | attaGGTCTCt***GGCT***GAGTTCAGAACAG | cloning THE1- and THE_G37D_-pt2 in pGGC000, "C" overhang for GreenGate system |
| THE1.pt2_BsaI_GGD_REV | taatGGTCTCt***CTGA***CCTTCCACGAGGATGAAC | cloning THE1- and THE_G37D_-pt2 in pGGC000, "D" overhang for GreenGate system |
| THE1_BsaI_GGC_FW2 | attaGGTCTCt***GGCT***ATGGTGTTCACAAAATCATTAC | cloning THE1_K636E_ in pGGC000 (incl. signal peptide), "C" overhang for GreenGate system |
| THE1_BsaI_GGD_REV2 | taatGGTCTCt***CTGA***CCTTCCACGAGGATGAACAAG | cloning THE1_K636E_ in pGGC000, "D" overhang for GreenGate system |
| SEQ_GG_FW | GTGAGTTAGCTCACTCATTAGGC | sequencing of inserts in pGGA000, pGGB000, pGGC000, pGGD000 |
| SEQ_GG_REV | GCTGCAAGGCGATTAAGTTGG | sequencing of inserts in pGGA000, pGGB000, pGGC000, pGGD001 |
| THE1_D634N_FW | CGTAATGTTAAAACCACGAATATCTTAC | site-directed mutagenesis to exchange D643 to N643 |
| THE1_D634N_REV | GTGTATAATGCTCTGCGATGCACC | site-directed mutagenesis to exchange D643 to N643 |
| THE1_Kinase_EcoRI_FW | attaGAATTCGTTGCTTCAAGGAAGCAGAGG | cloning the kinase domains of THE1_K636E_ and THE1_D634N_ in pGEX-6P-1 |
| THE1_Kinase_STOP_NotI_REV | taatGCGGCCGCTTACCTTCCACGAGGATGAACAAG | cloning the kinase domains of THE1_K636E_ and THE1_D634N_ in pGEX-6P-1 |
| THE1pro_seq_FW | GCTTCAAACTTTCCTCTTCTCTAC | amplifying THE1_pro_-THE1-GFP sequence from THE1-GFP or THE1_G37D_-GFP-transformed plants for verifying its integrity by sequencing |
| eGFP_seq_REV | GTCTTGTAGTTGCCGTCGTC | amplifying THE1pro-THE1-GFP sequence from THE1-GFP or THE1_G37D_-GFP-transformed plants for verifying its integrity by sequencing |

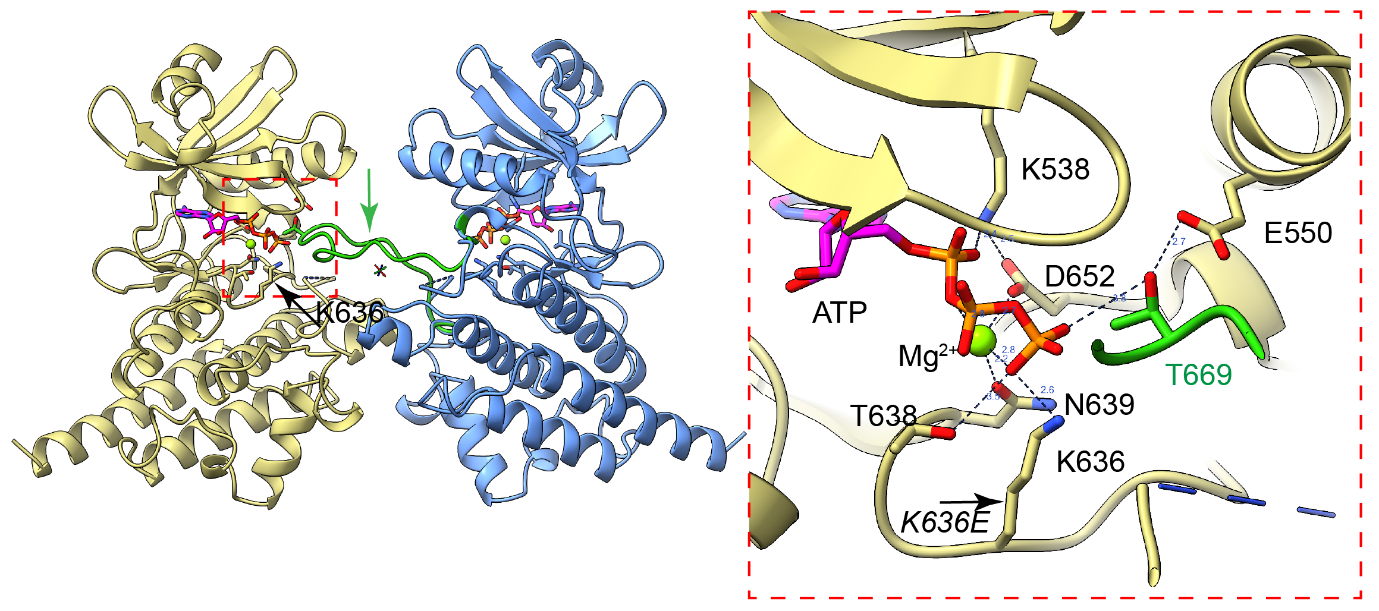

**Figure S1** Structural model of a THESEUS1 kinase-domain dimer with the activation loop of one protomer bound in the active site of the other. *(Left)* Overview of the modelled THE1 kinase-domain dimer (protomers in yellow and blue). The activation loops are shown in green; the arrow marks the segment of the blue protomer’s activation loop that reaches into the active site of the yellow protomer. *(Right)* Close-up of the boxed region. ATP (magenta/orange) and Mg^2+^ (green sphere) are shown as sticks and sphere; dashed lines indicate polar contacts, distances in Å.

The model is based on the AlphaFold prediction for *A. thaliana* THE1 (UniProt Q9LK35; kinase domain, residues 493–795). The kinase fold and the position of the nucleotide were taken from the crystal structure of the FERONIA kinase domain (PDB 7XDV, 1.97 Å; UniProt Q9SCZ4), onto which the THE1 model was superposed, and ATP·Mg^2+^ was placed accordingly. The arrangement of the two protomers was generated by manual docking and subsequently regularized by interactive molecular dynamics flexible fitting (ISOLDE) and real-space refinement (Coot). This dimer arrangement is a model and is not observed in the 7XDV crystal lattice.

The active site contains the canonical protein-kinase catalytic machinery: K538 (β3 lysine) and E550 (αC glutamate) position the phosphates of ATP; N639 and D652 (DFG motif) coordinate the Mg^2+^ ion; and D634 of the catalytic HRD motif (H632-R633-D634) acts as the catalytic base that deprotonates the acceptor hydroxyl. K636 is the catalytic-loop lysine of the HRD-x-K motif (H632-R633-D634-V635-K636-T637-T638-N639) and contacts the γ-phosphate of ATP directly. This lysine is invariant in Ser/Thr protein kinases — K168 in PKA and K663 in FERONIA, where it is resolved in 7XDV. Its positive charge neutralizes the developing negative charge on the transferred phosphoryl group and thereby stabilizes the transition state of phosphoryl transfer. T669, a reported *in vivo* phosphosite of the THE1 activation loop (Durek et al. 2010), is presented *in trans* by the partner protomer and points towards the γ-phosphate, consistent with trans-autophosphorylation of the activation loop.

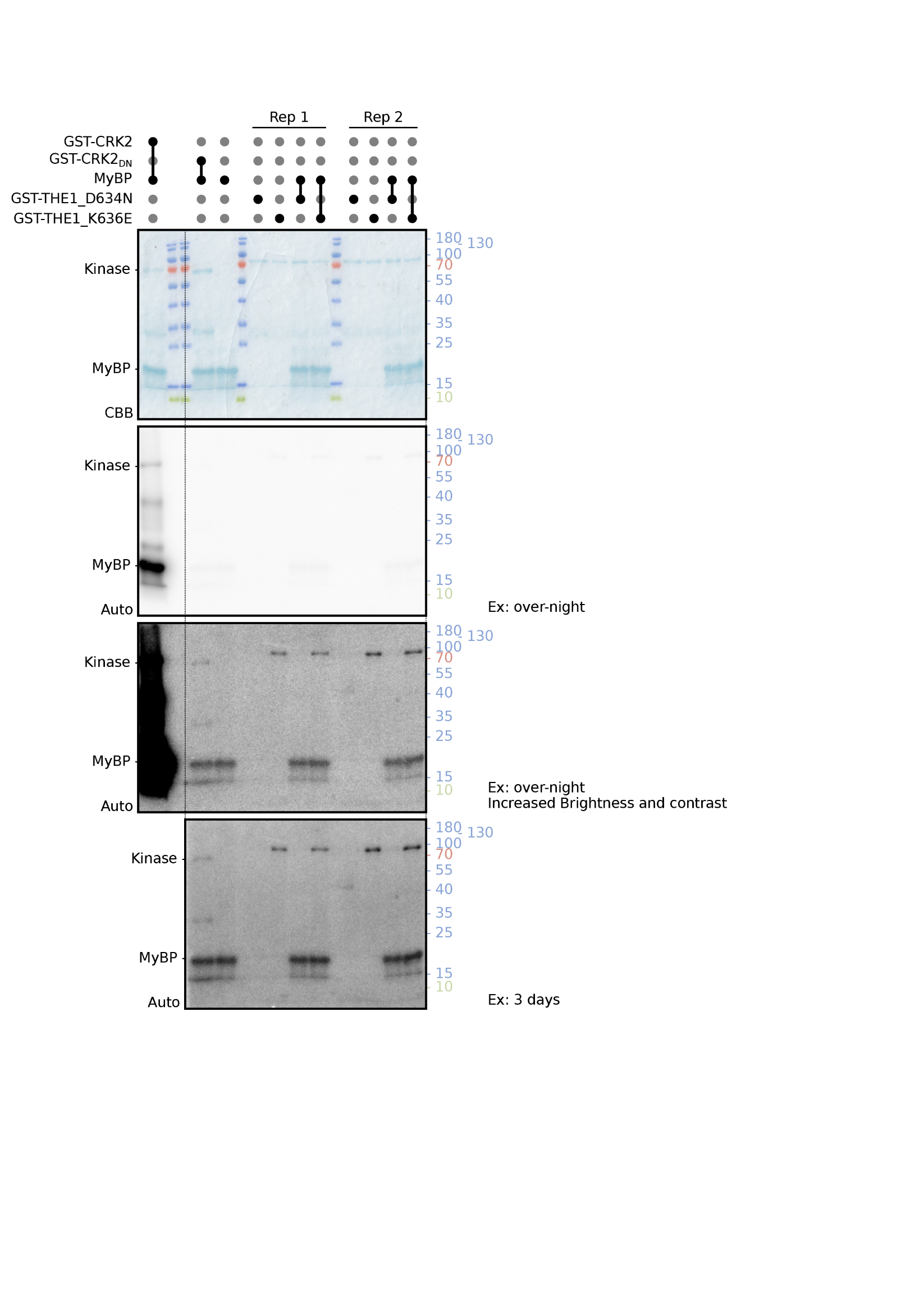

**Figure S2** Kinase assay showing activity of THE1 mutated variants. GST-THE1_K363E_ shows weak autophosphorylation activity, while GST-THE1_D634N_ shows no detectable activity. No THE1 transphosphorylation of myelin basic protein (MyBP) was observed, in comparison to GST-CRK2. GST-CRK2 and its kinase dead variant GST-CRK2_DN_ were used as controls. CBB – Coomassie brilliant blue, Auto – autoradiography, Rep - repeat, Ex - exposure. Marker in kDa.

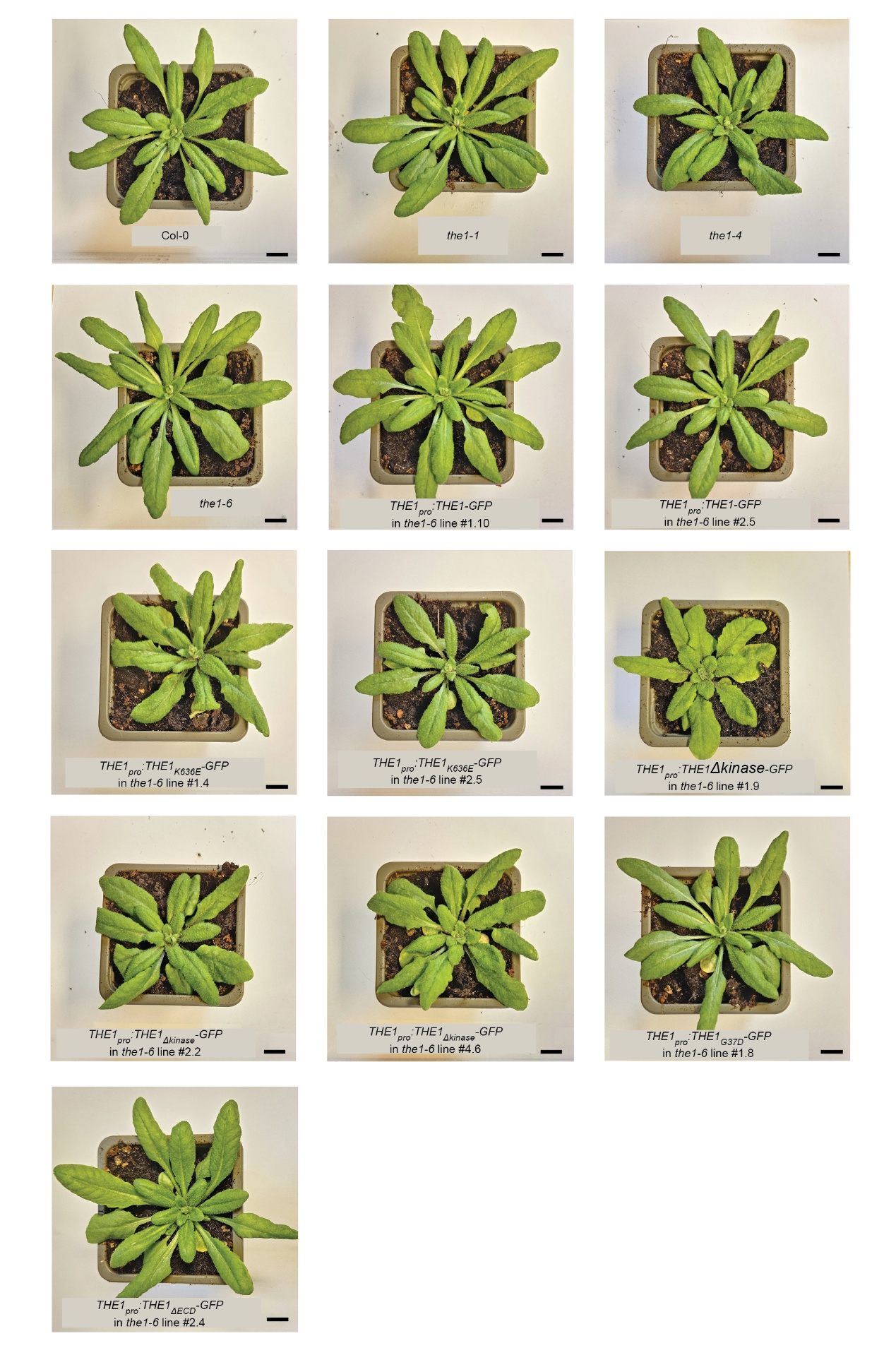

**Figure S3** Representative images of rosettes of four-week-old *A. thaliana* wildtype plants, *THE1* mutants, and *THE1-GFP* variant transgenic lines. Plants were soil-grown in 16 hour light/8 hour dark conditions in a shared tray with pots in a randomized location. Scale bar: 1 cm.

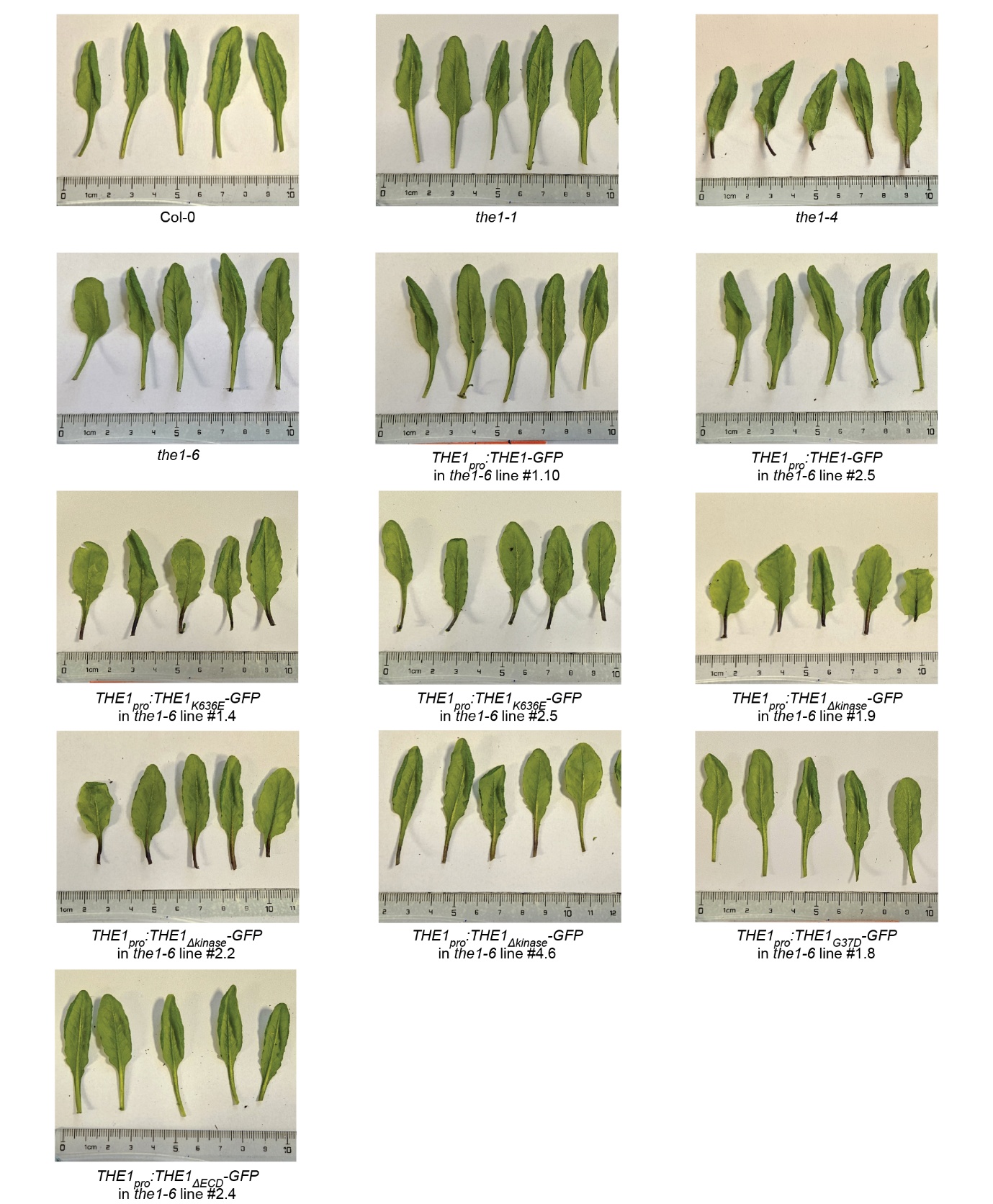

**Figure S4** Representative images of five true leaves starting from the 6^th^ true leaf in rosettes of four-week-old *A. thaliana* wildtype plants, *THE1* mutants, and *THE1-GFP* transgenic lines. All leaves shown for a line come from the same plant. Plants were soil-grown in 16 hour light/ 8 hour dark conditions in a shared tray with pots in a randomized location.

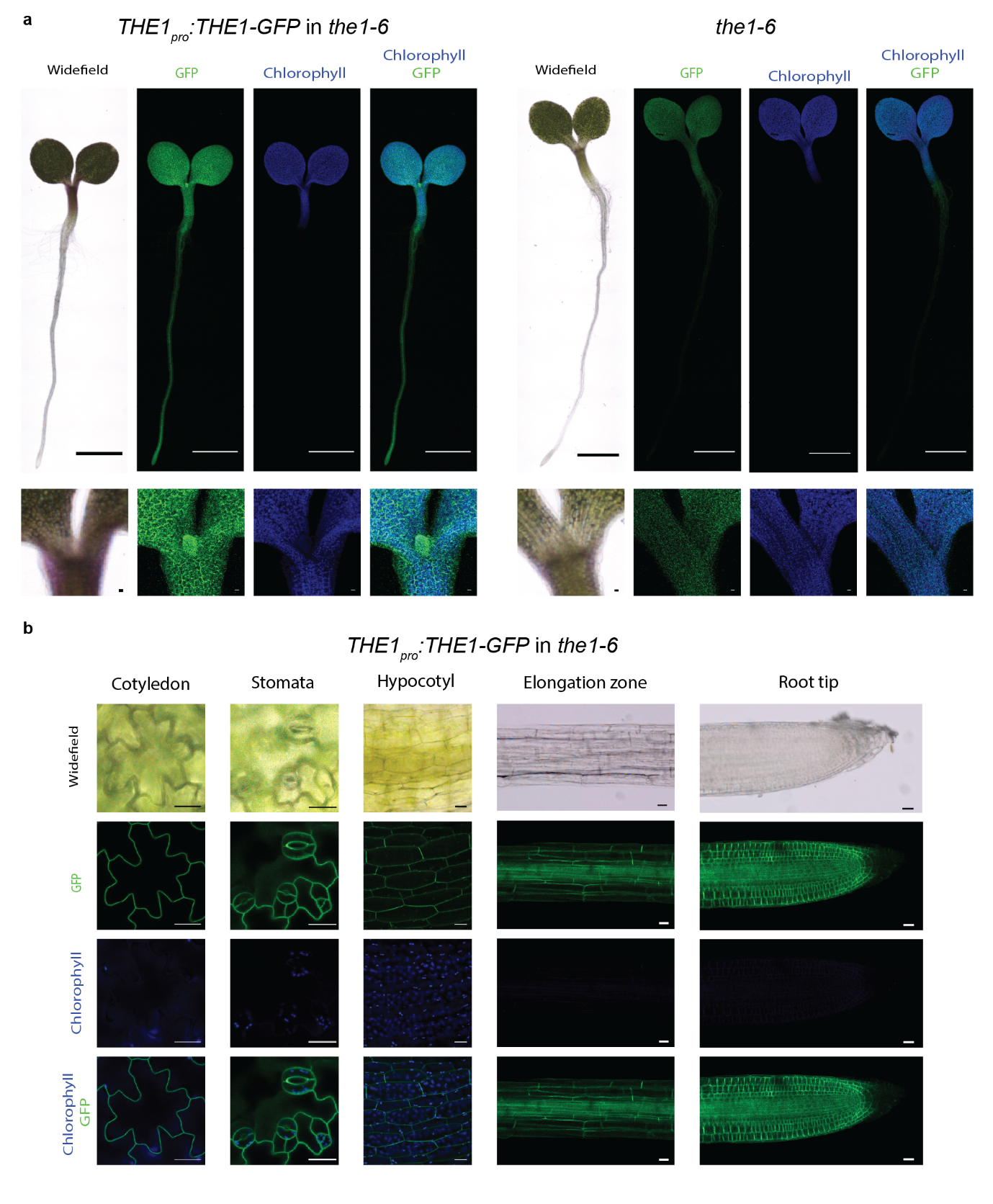

**Figure S5** Overview of *THE1_pro_:THE1-GFP* expression in a six-day-old *A. thaliana* seedling. **a)** Comparison of a *the1-6* seedling transformed with *THE1_pro_:THE1-GFP* to an untransformed control. **b)** Close-ups of the cotyledon epidermis (adaxial side), stomata, hypocotyl epidermis, elongation zone and root tip. Scale bar: 1 mm (seedling) or 20 µm (other images).

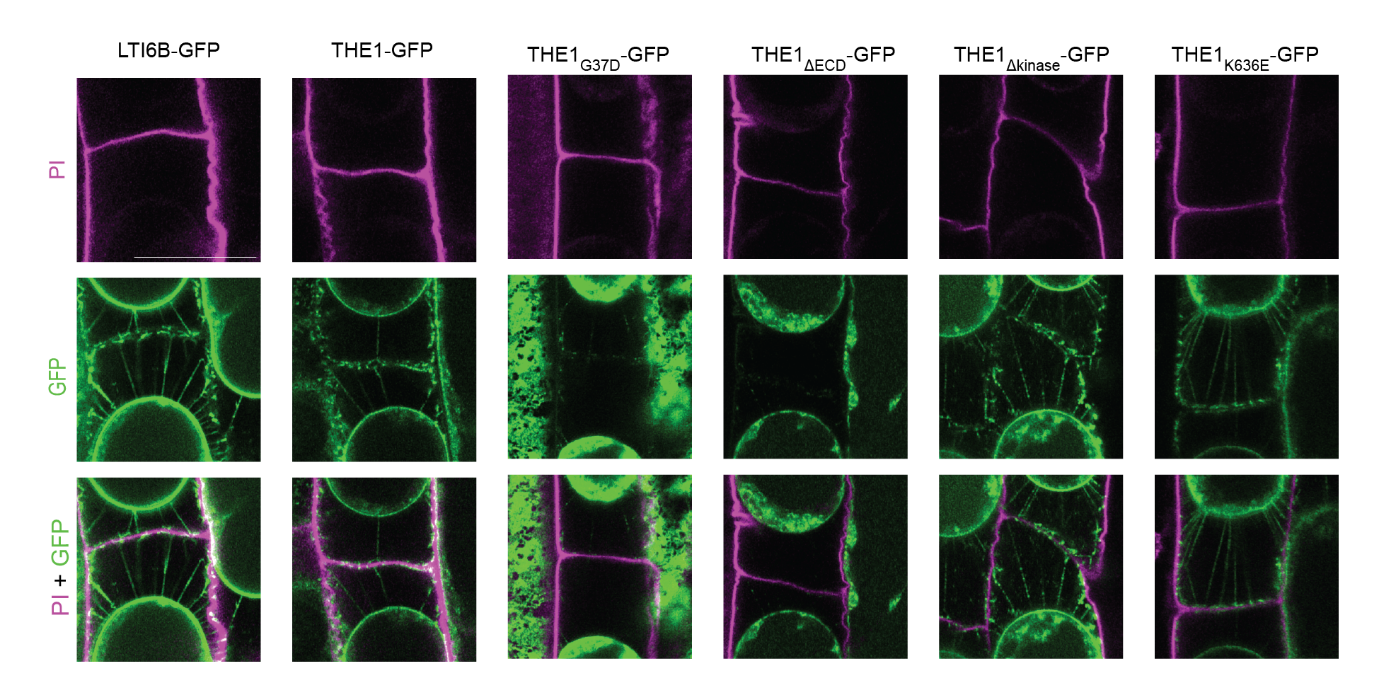

**Figure S6** Plasmolyzed epidermal cells in the elongation zone of six-day-old *A. thaliana* seedlings expressing the plasma membrane marker LTI6B-GFP or THE1-GFP variants. Plants were plasmolyzed in 450 mM sorbitol, 15 µM propidium iodide (PI), ½ MS for 15 minutes. Scalebar: 20 µm.

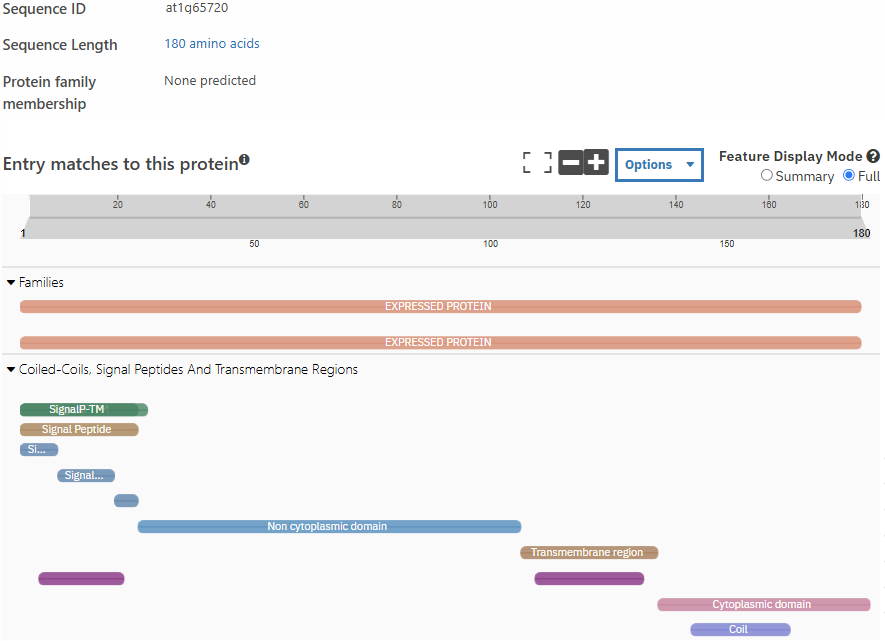
**Figure S7** InterProScan annotation of the *A. thaliana* protein of unknown function AT1G65720 (Blum et al. 2026). The full-length amino acid sequence of AT1G65720 was used as query sequence and the query was submitted on the 3^rd^ of September 2026.

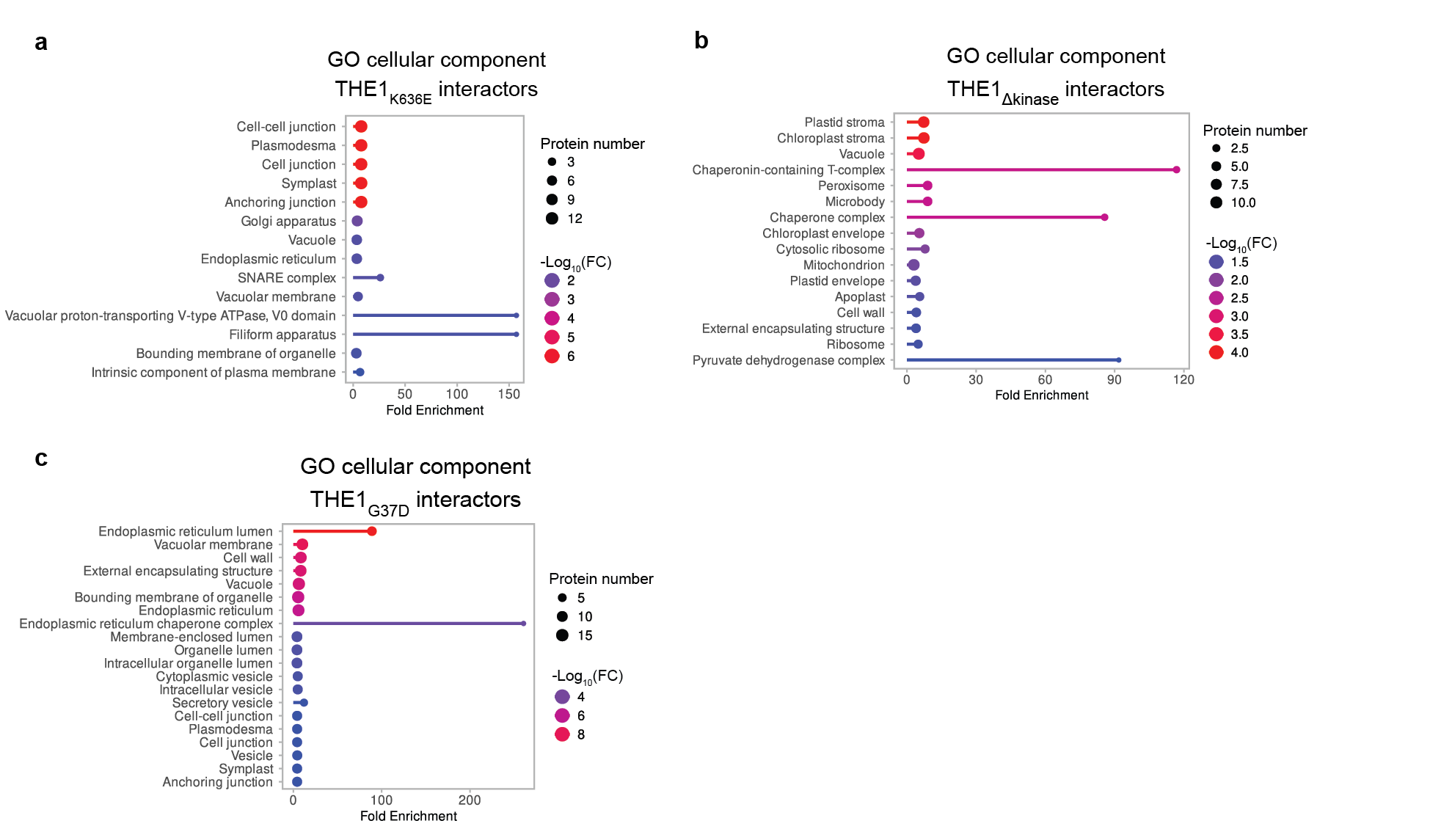

**Figure S8** Significantly enriched cellular component gene ontology (GO)-terms associated with THE1 interactors. **a)** THE1_K636E_. **b)** THE1_Δkinase_. **c)** THE1_G37D_. Circle size: number of interactors associated with the GO term. Circle color: ‑log_10_ of false discovery rate (FDR)-adjusted p-values. FDR cutoff for GO enrichment analysis: p-value <0.05.

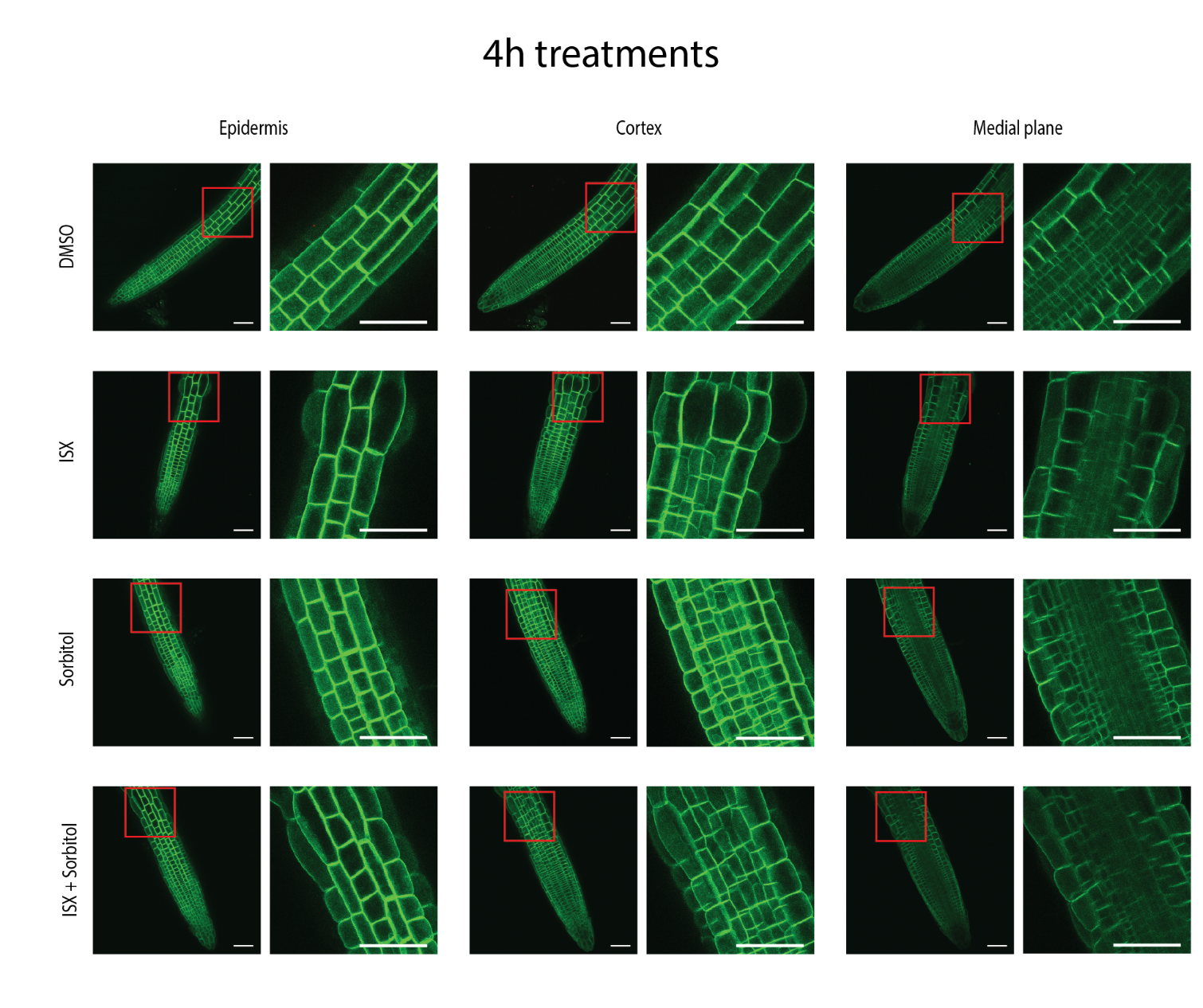

**Figure S9** Localization of THE1-GFP in six-day-old *A. thaliana* seedling roots expressing *THE1_pro_:THE1-GFP* after four-hour treatment with DMSO (solvent control), ISX (600 nM), sorbitol (300 mM), or combined ISX-sorbitol (600 nM and 300 mM respectively).

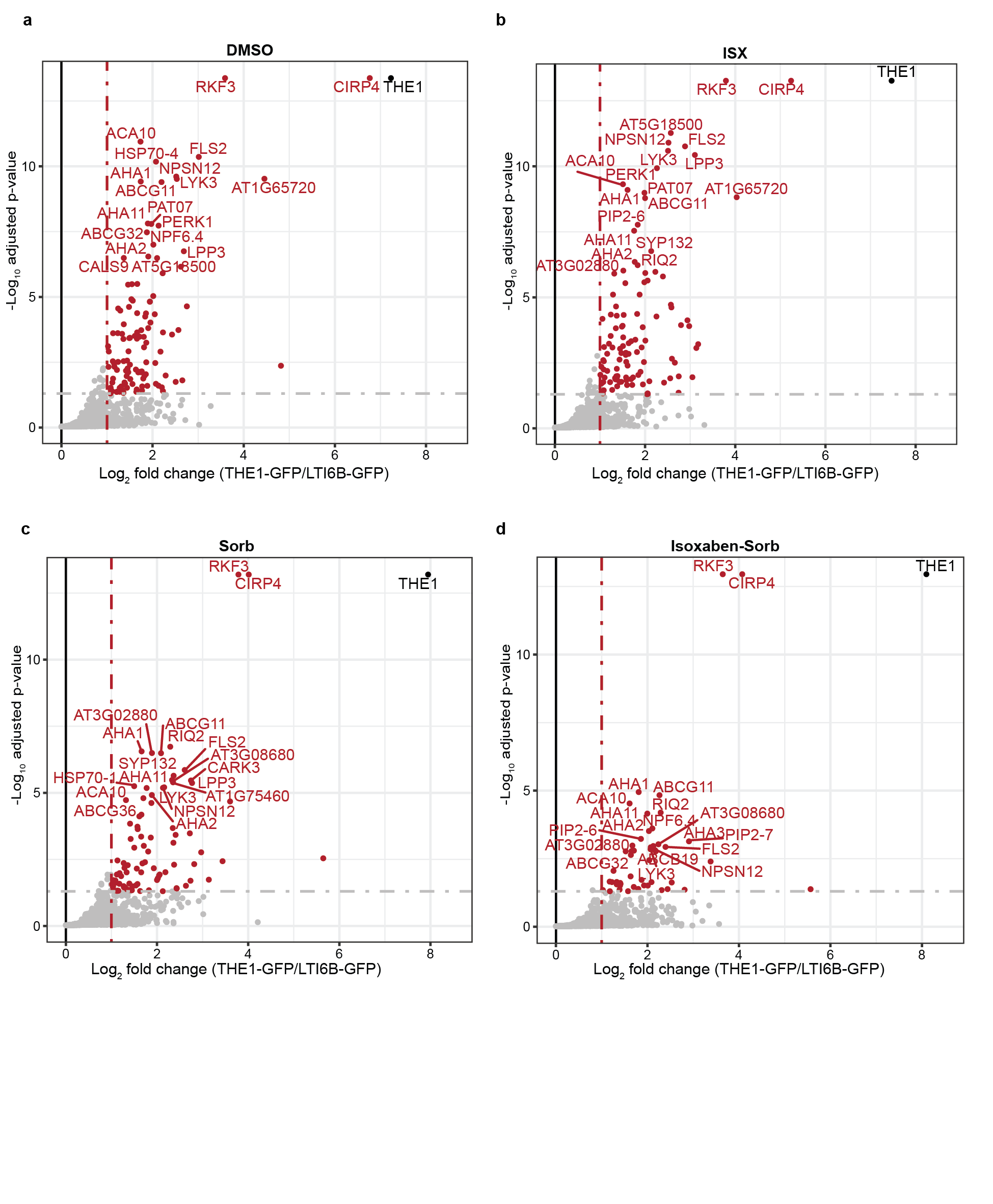

**Figure S10** Putative THE1 interactors identified by immunoprecipitation–liquid chromatography–tandem mass spectrometry in seven-day-old *A. thaliana* seedlings expressing *THE1_pro_:THE1-GFP* after four-hour treatment with: **a)** DMSO (solvent control), **b)** ISX (600 nM), **c)** sorbitol (300 mM), or **d)** combined ISX-sorbitol (600 nM and 300 mM respectively) treatment (ISX-Sorb). Putative interactors were identified by enrichment analysis with the R-package *limma* (Ritchie et al. 2015) and false discovery rate (FDR) correction using the Strimmer’s Q-value method. Proteins with a fold-change (FC) >2 and adjusted P-value <0.05 were considered significantly enriched. Red dots: proteins identified as significantly enriched interactors; grey dots: non-significant enriched proteins; black dot: the THE1 bait protein.

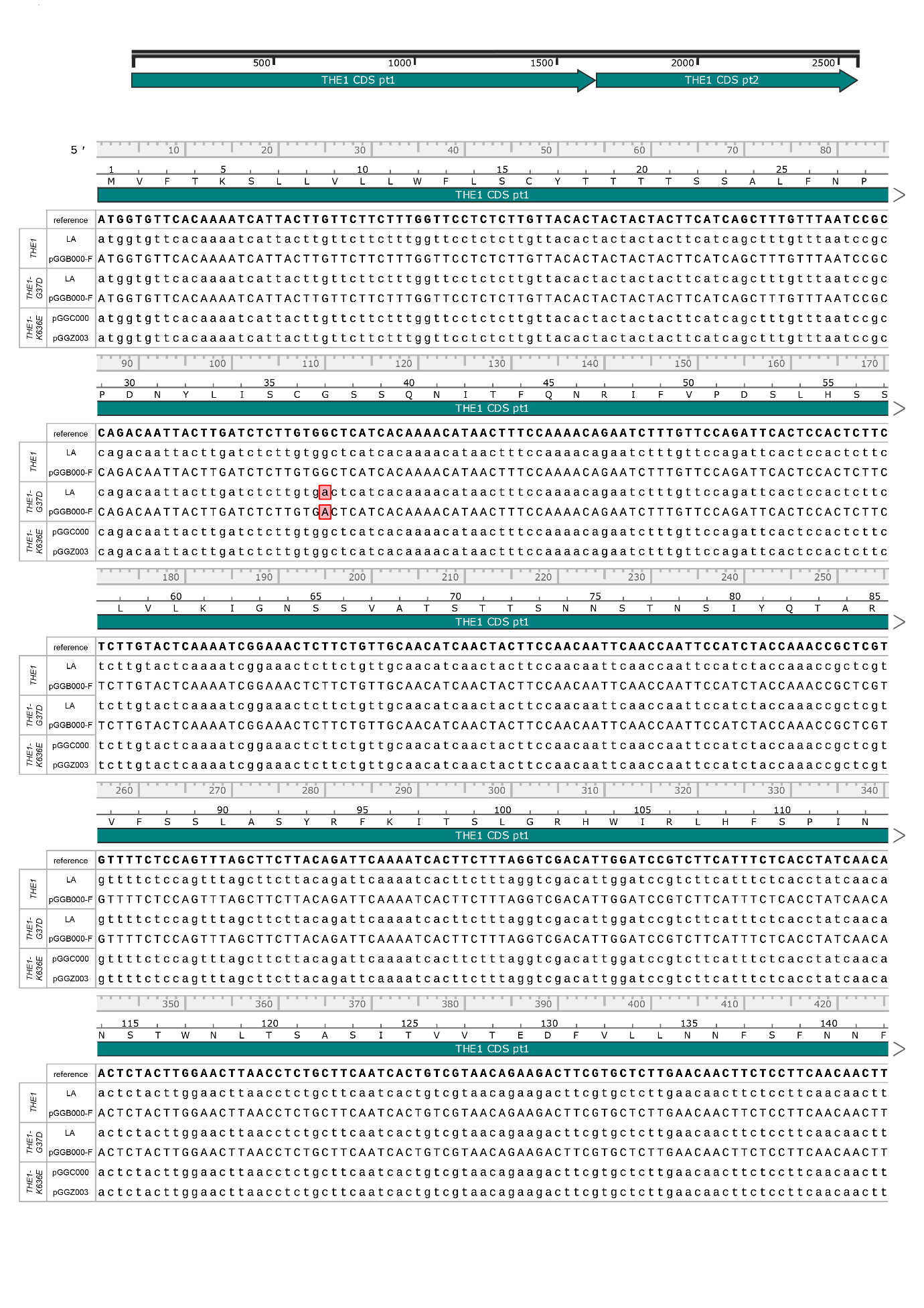

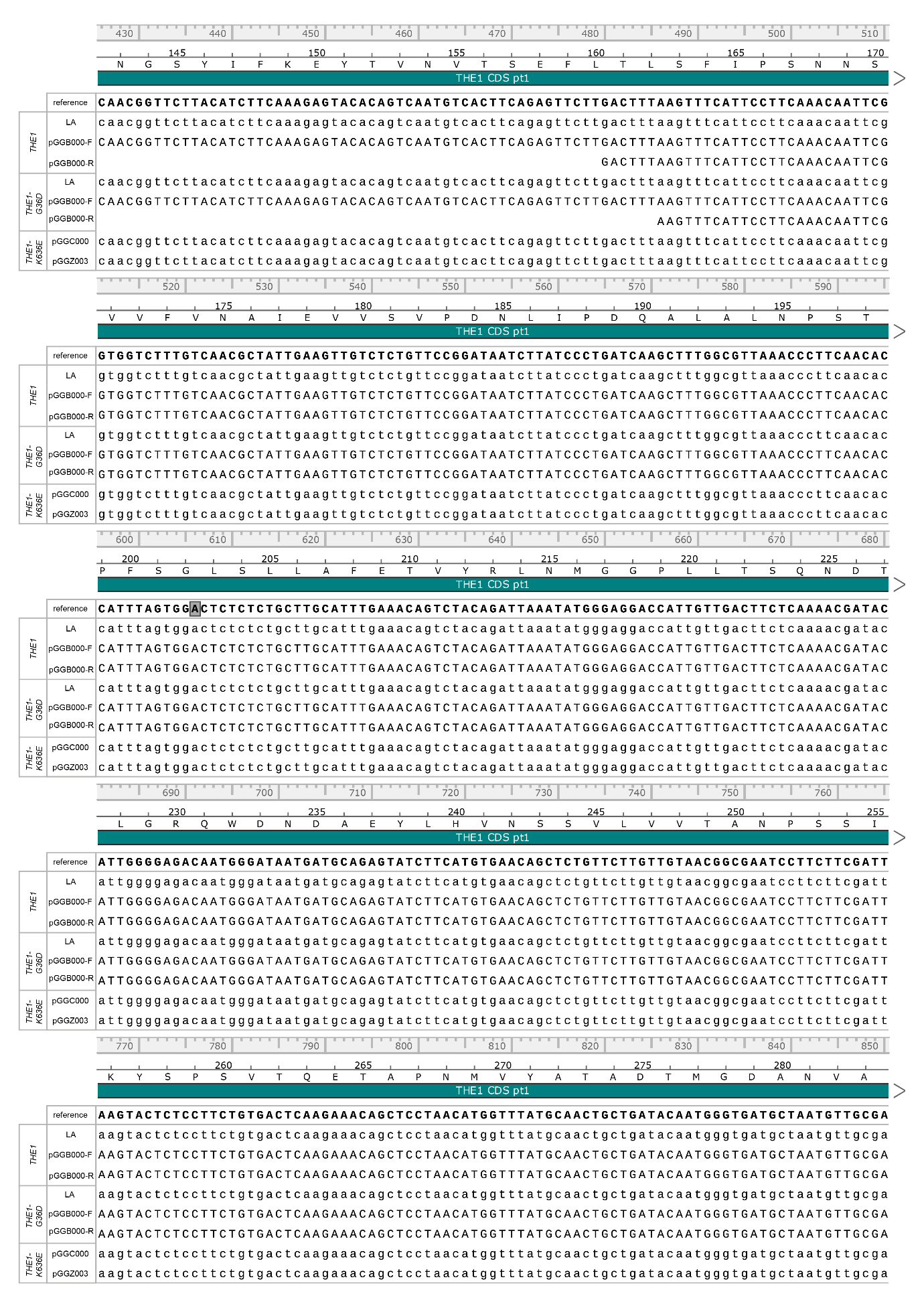

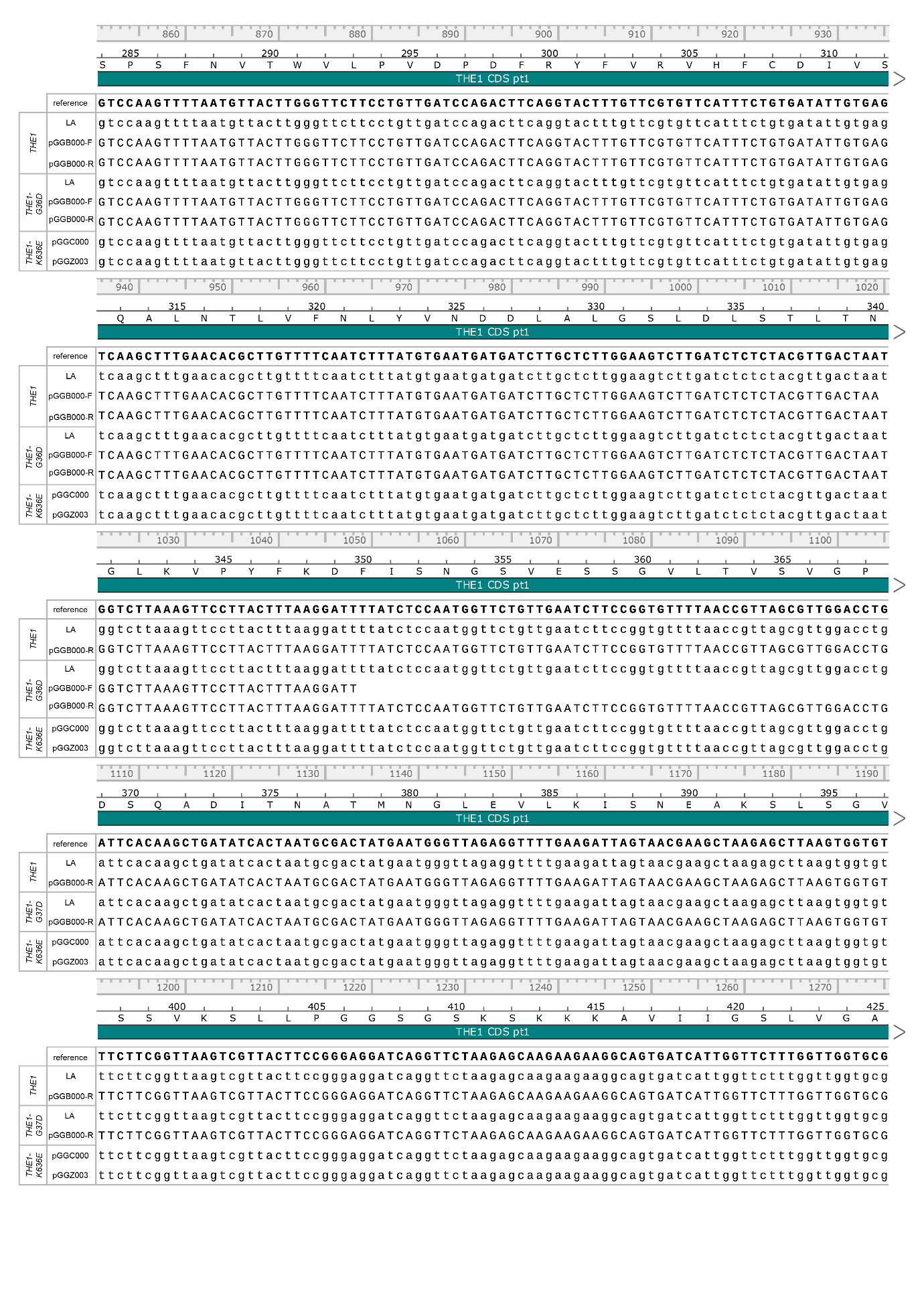

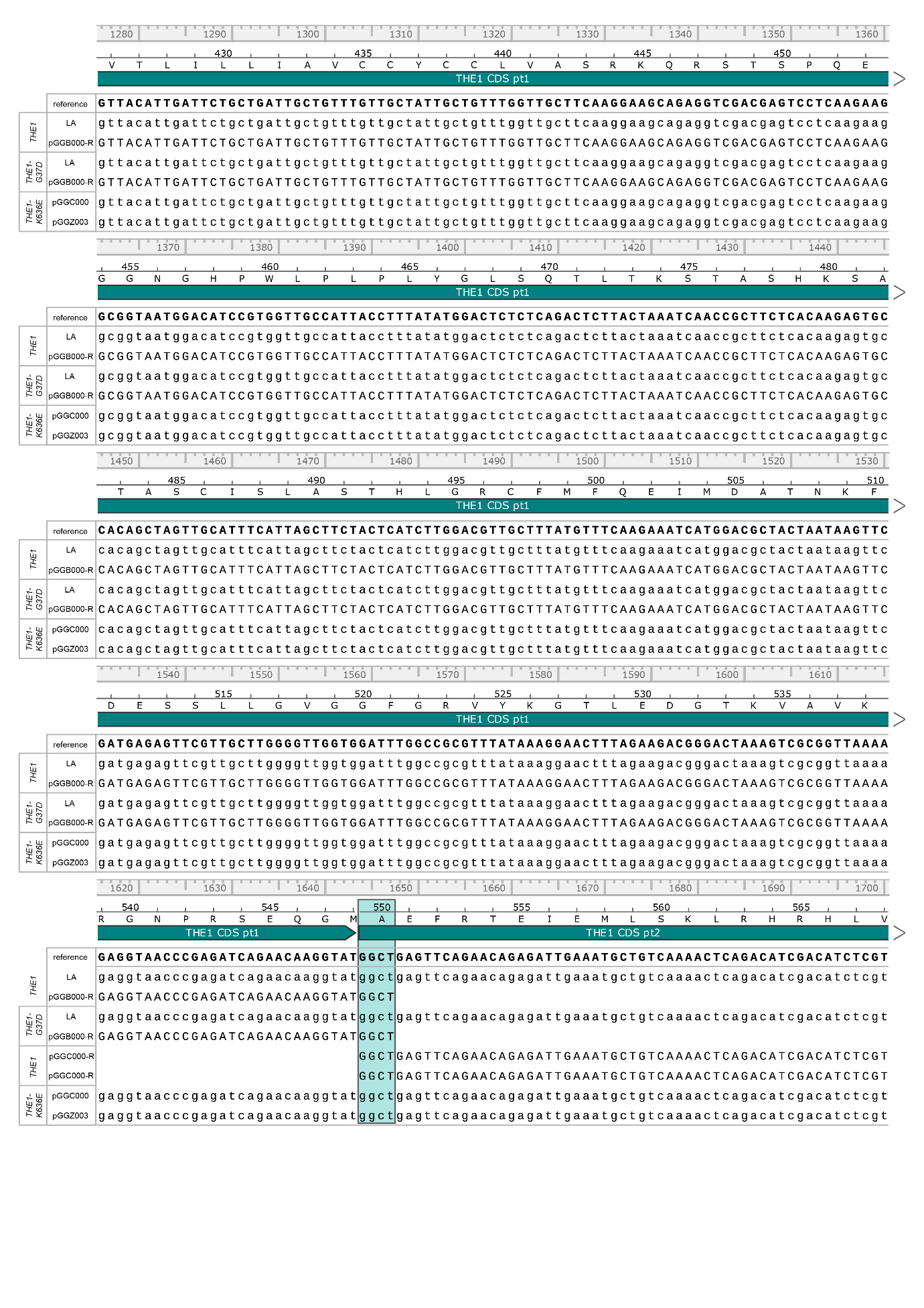

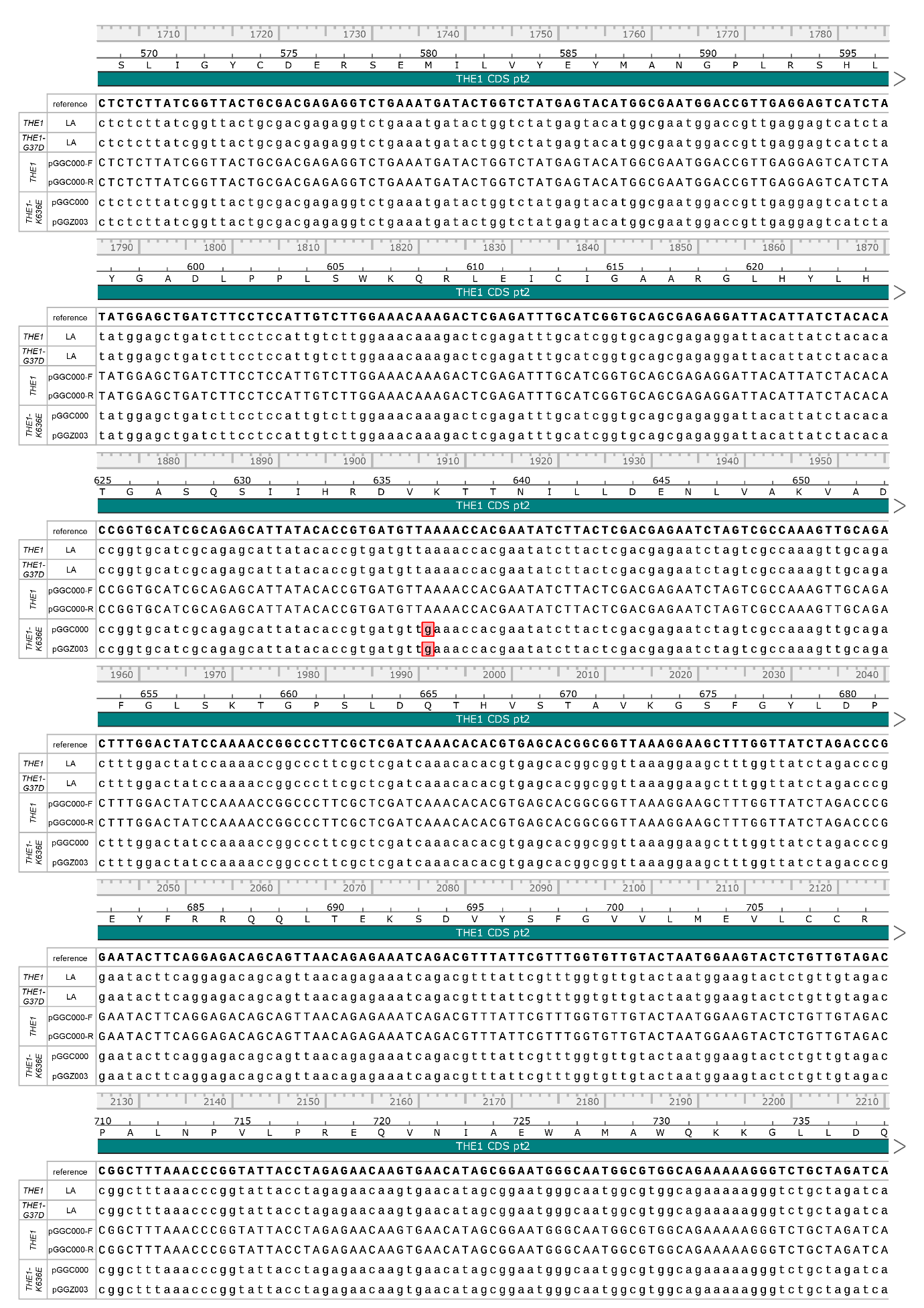

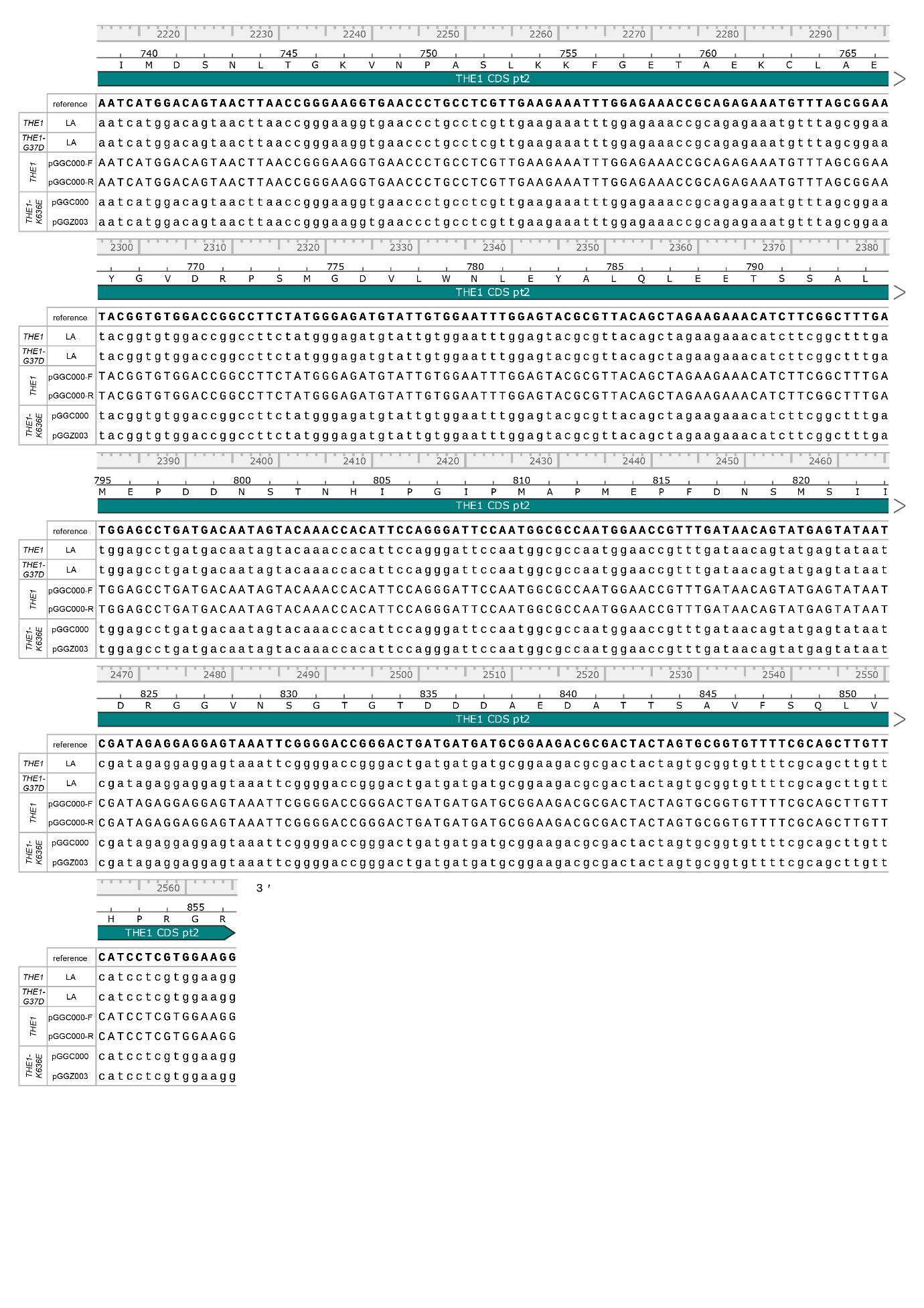

**Figure S11**: Sequencing of *THE1* constructs. Linear amplicon (LA) results are sequencing results of *THE1* and *THE1_G37D_* as amplified from *Agrobacterium tumefaciens* containing pGGZ003 with *THE1-GFP* or *THE1_G37D_-GFP*. The plasmid on which the results from the other sequences are located is indicated. Lowercase sequences for which the reference is not LA are whole plasmid Oxford Nanopore Technology long-read results. Uppercase sequences are Sanger sequencing-based results. The 110g>a mutation in *THE1_G37D_* and 1906a>g mutation in *THE1_K636E_* are indicated in red. We removed the *Bsa*I site in *THE1* by changing nucleotide position 606 (indicated with grey box) from T to A. The full-length THE1 sequence could not be cloned in *E. coli*. We cloned THE1 in two parts split on the GGCT B-C overhang (THE1 nucleotides 1644-1647, indicated in blue) of the GreenGate cloning system, used these parts in the assembly of the final transformation vector, and transformed this directly into *A. tumefaciens*.

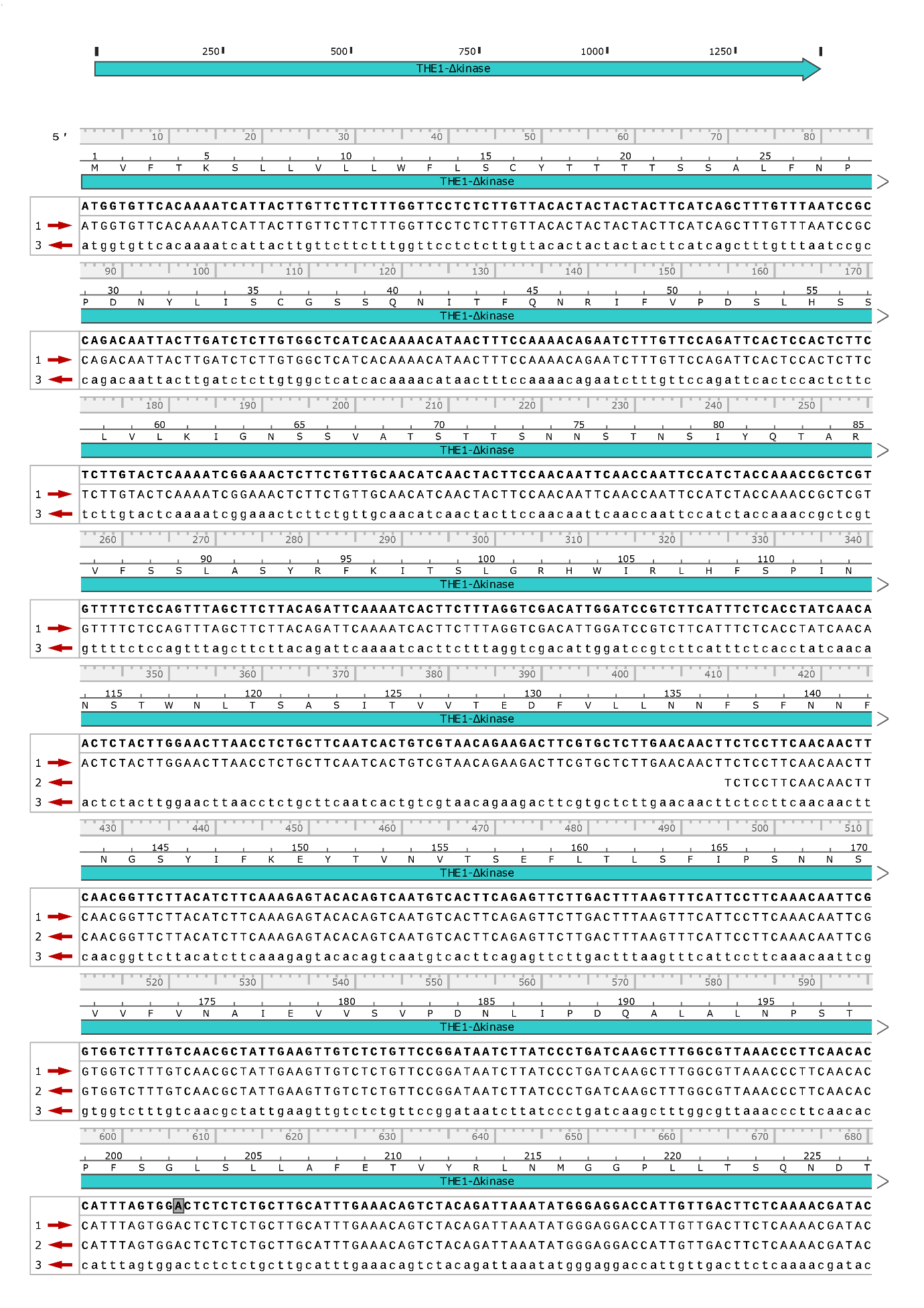

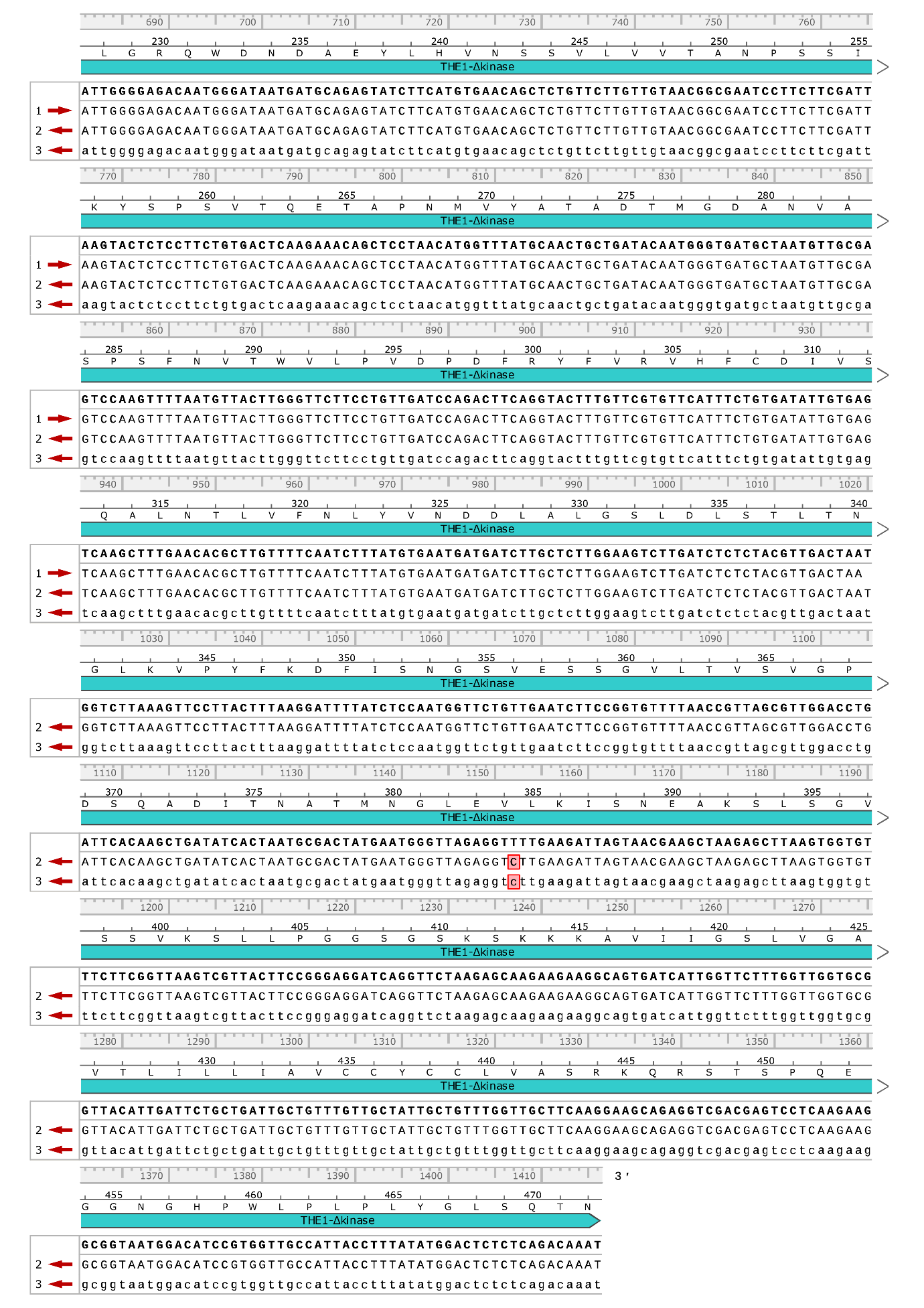

**Figure S12**: Sanger sequencing results of *THE1_Δkinase_* in pGGC000 (1 + 2) and Oxford Nanopore Technologies sequencing results of *THE1_Δkinase_* in pGGZ003 (3). The reference sequence is shown in bold. The *Bsa*I site in *THE1* was removed by changing nucleotide position 606 (indicated with grey box) from T to A. There is a synonymous mutation in nucleotide position 1152, where a T mutated to a C, the amino acid remains a Valine.

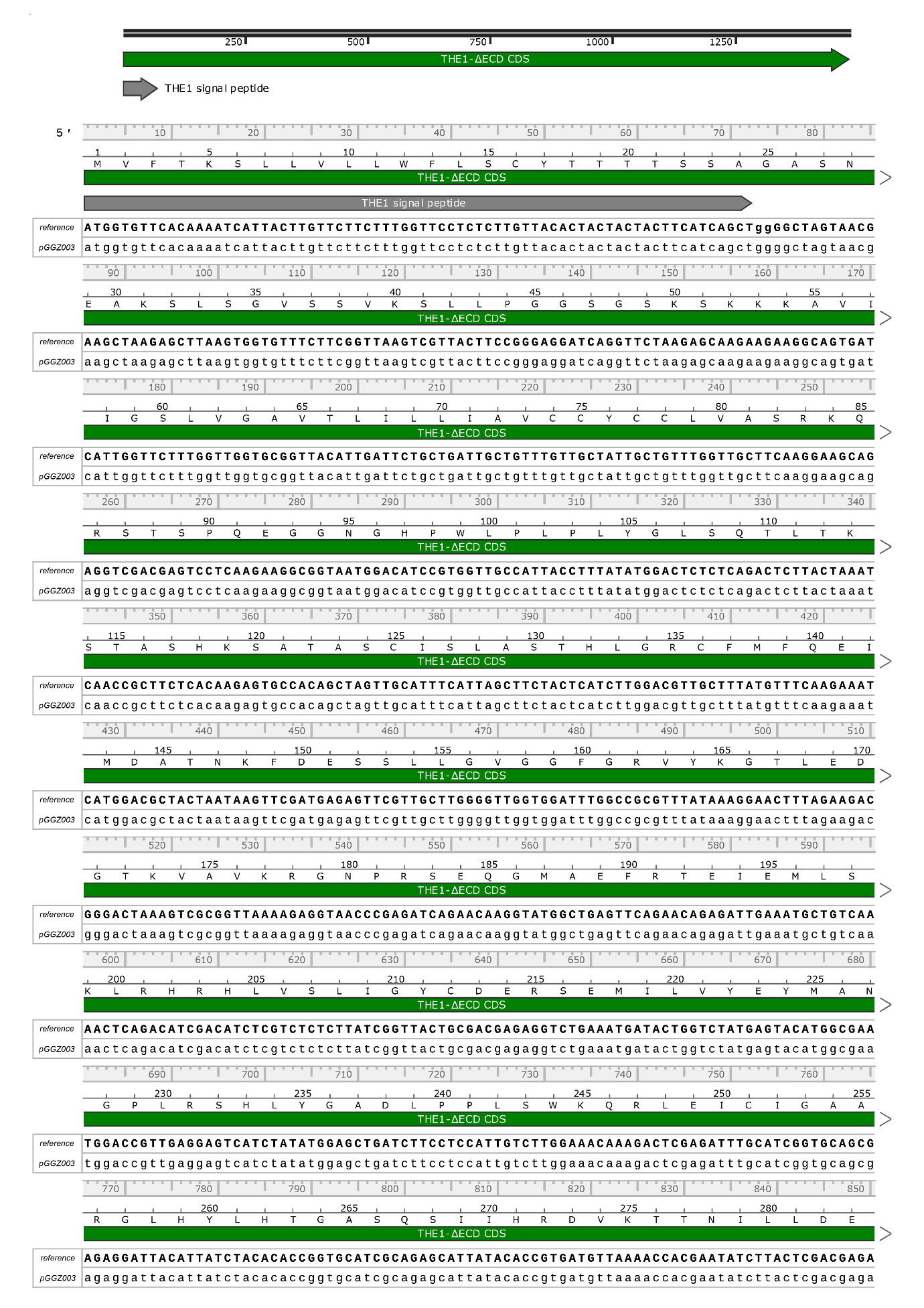

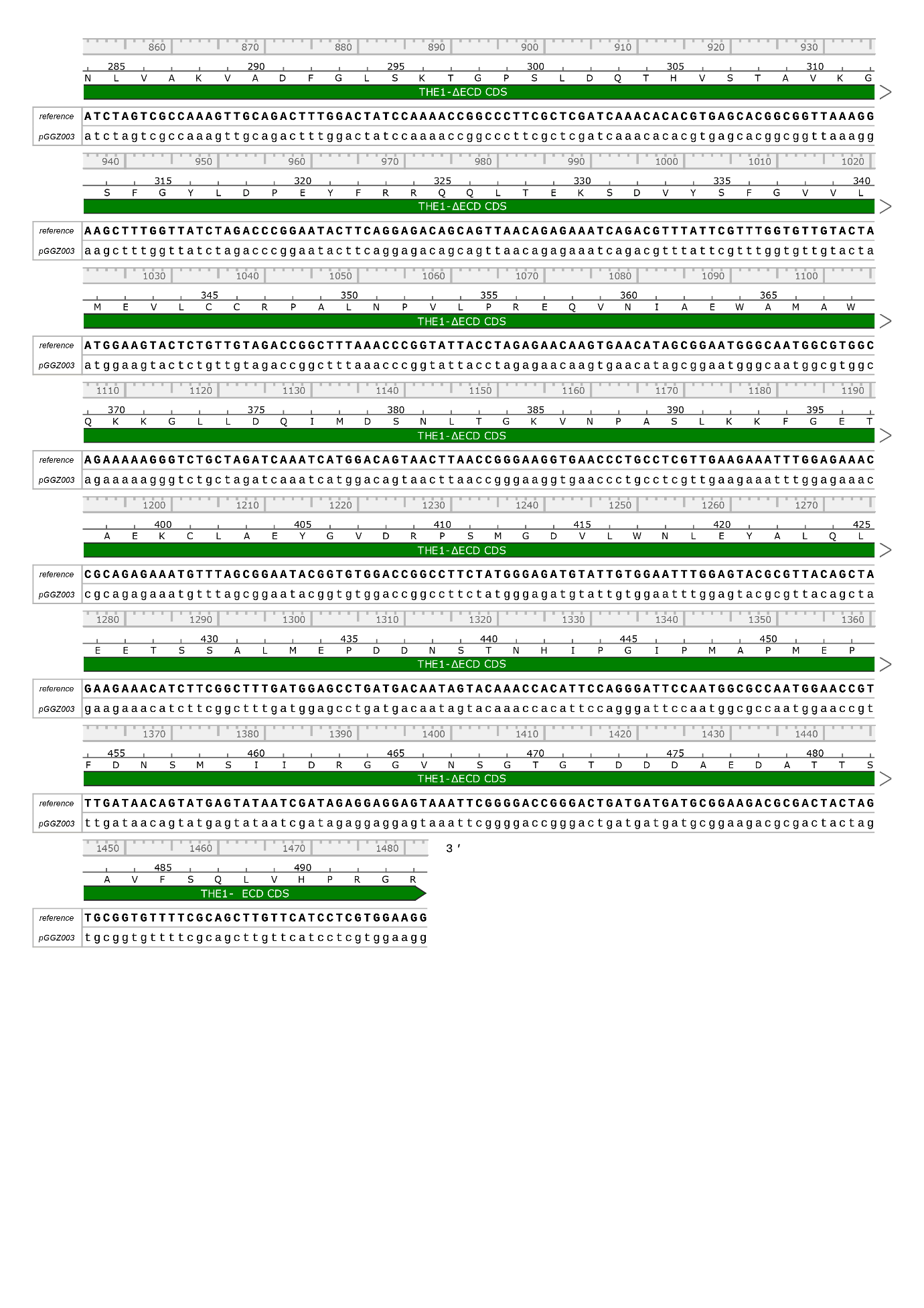

**Figure S13**: Oxford Nanopore Technologies long-read sequencing result of *THE1_ΔECD_* in the pGGZ003 destination plasmid used for plant transformations to obtain transgenic lines containing *THE1_ΔECD_-GFP*.

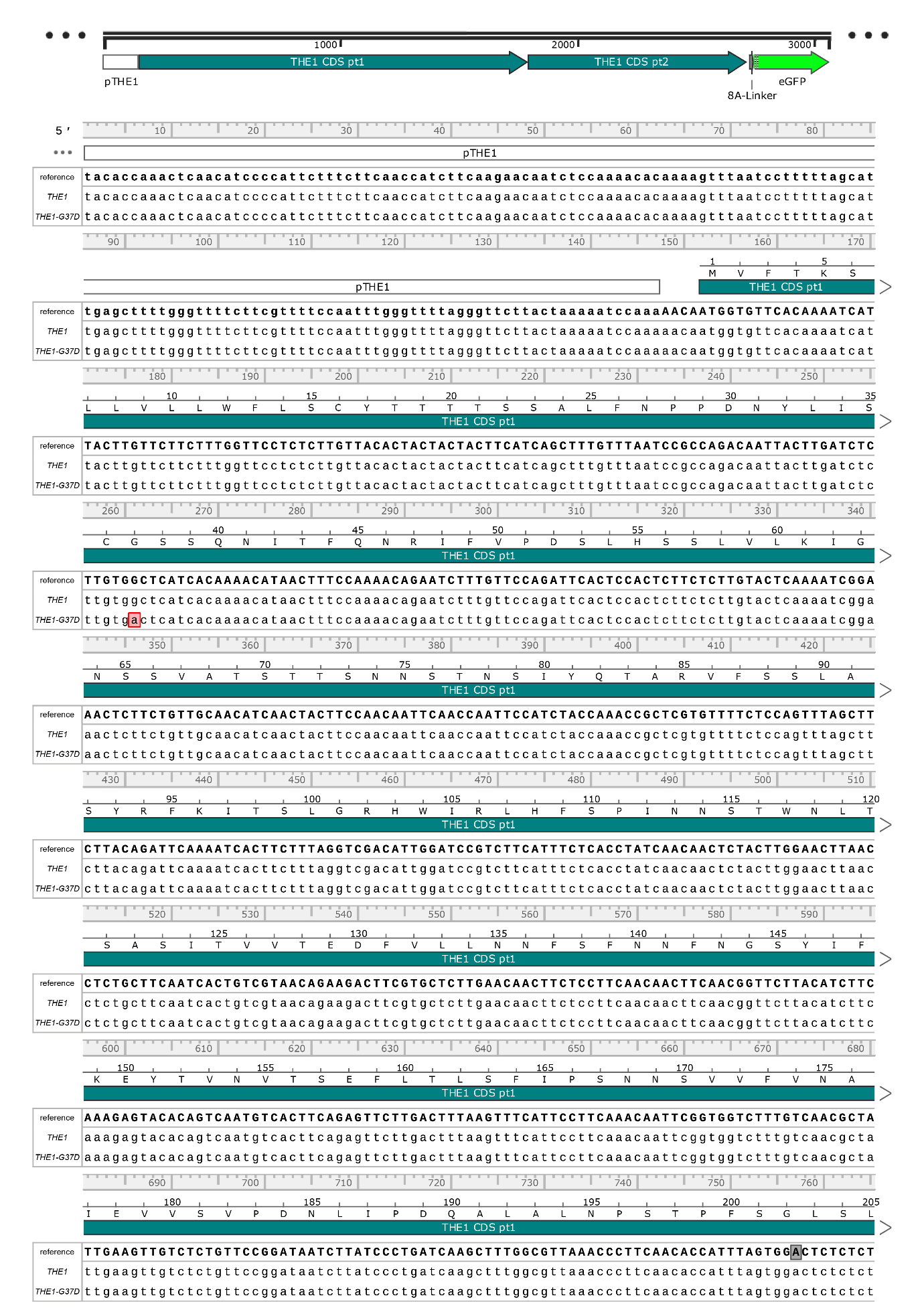

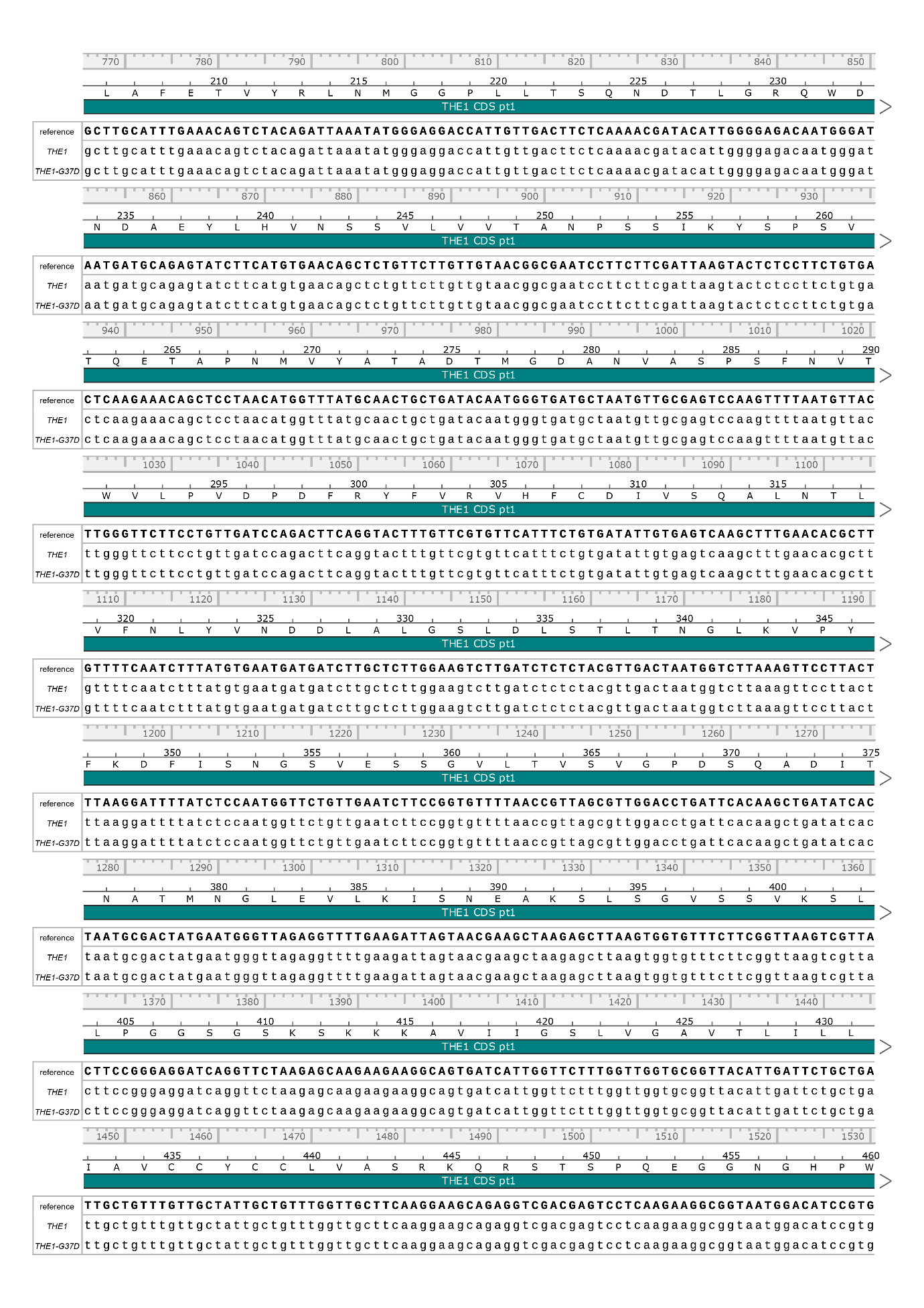

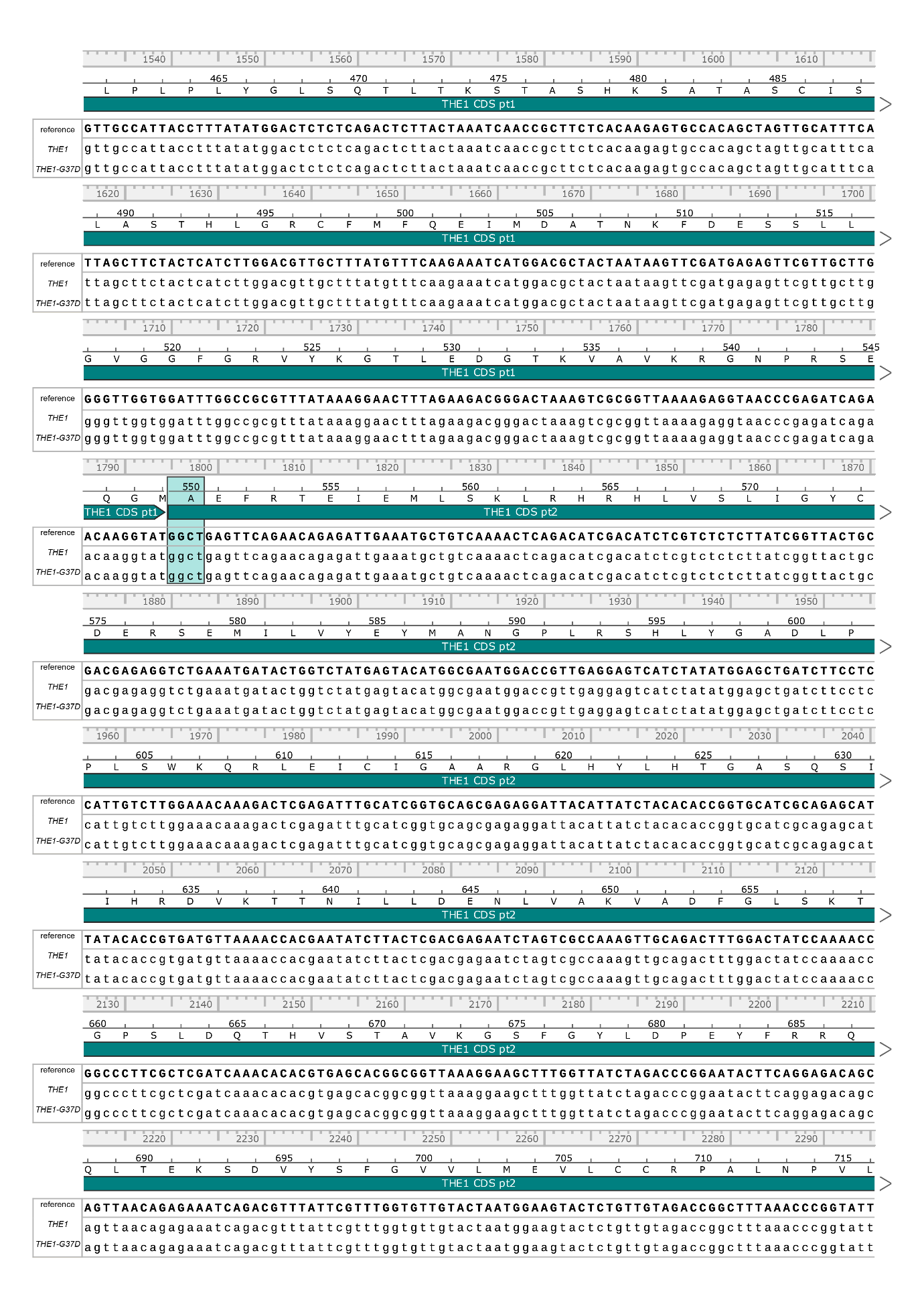

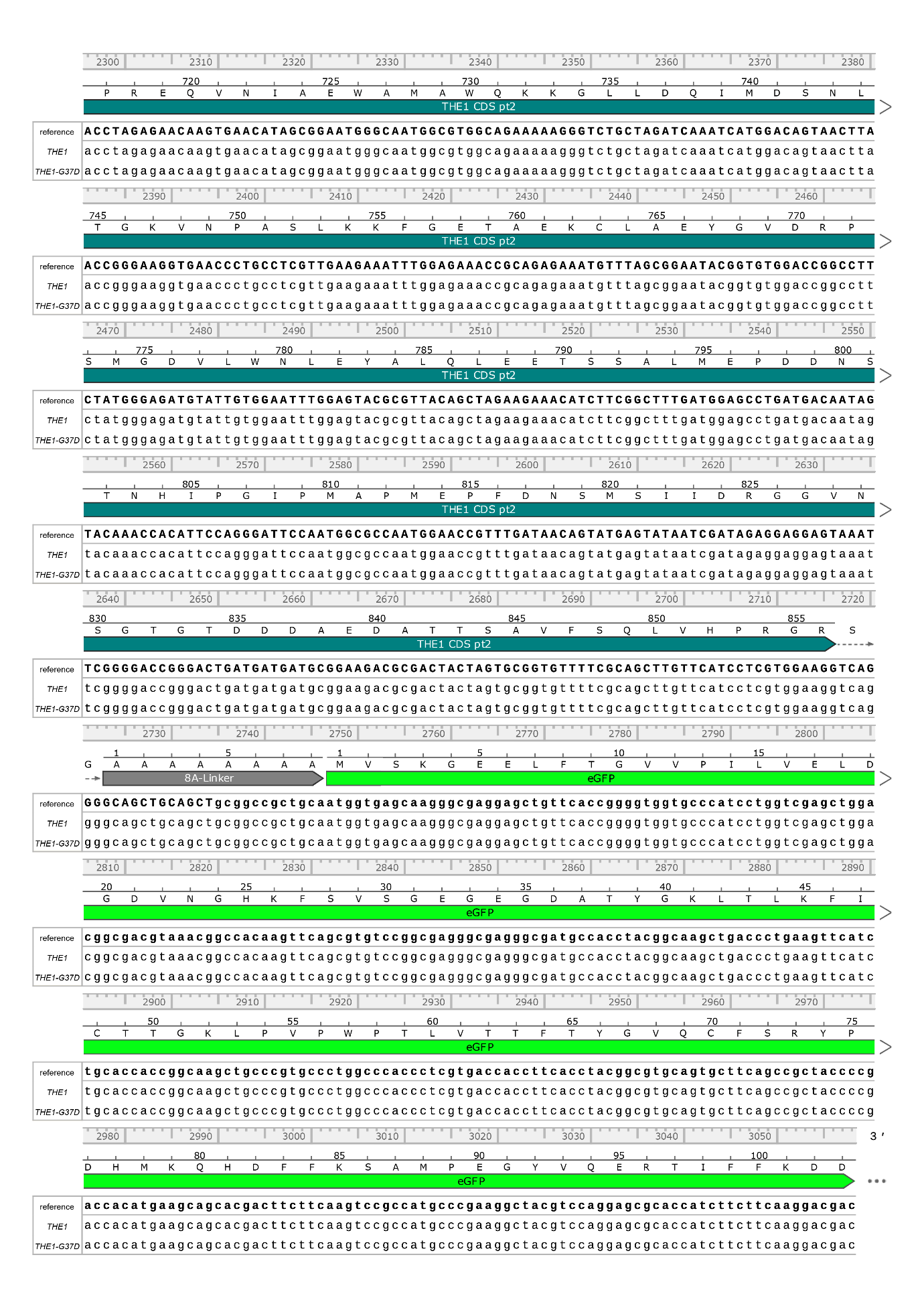

**Figure S14**: Oxford Nanopore Technologies long-read LA sequencing results of *THE1-GFP* and *THE1_G37D_-GFP* amplified from transgenic *A. thaliana* lines used in this study (*THE1_pro_:THE1-GFP* #1.10 and *THE1_pro_:THE1_G37D_-GFP* #1.8). The 110g>a mutation in *THE1_G37D_* is indicated in red. The *Bsa*I site in the *THE1* sequence was removed by changing nucleotide position 606 (indicated with a grey box) from T to A.
